# Bats dynamically change echolocation parameters in response to acoustic playback

**DOI:** 10.1101/604603

**Authors:** M. Jerome Beetz, Manfred Kössl, Julio C. Hechavarría

## Abstract

Animals extract behaviorally relevant signals from “noisy” environments. To investigate signal extraction, echolocating provides a rich system testbed. For orientation, bats broadcast calls and assign each echo to the corresponding call. When orienting in acoustically enriched environments or when approaching targets, bats change their spectro-temporal call design. Thus, to assess call adjustments that are exclusively meant to facilitate signal extraction in “noisy” environments, it is necessary to control for distance-dependent call changes. By swinging bats in a pendulum, we tested the influence of acoustic playback on the echolocation behavior of *Carollia perspicillata*. This paradigm evokes reproducible orientation behavior and allows a precise definition of the influence of the acoustic context. Our results show that bats dynamically switch between different adaptations to cope with sound-based navigation in acoustically contaminated environments. These dynamics of echolocation behavior may explain the large variety of adaptations that have been reported in the bat literature.

**Summary statement:** The frugivorous bat *Carollia perspicillata* dynamically switch between different adaptations when echolocating in acoustically contaminated environments.

## Introduction

For orientation, echolocating bats emit biosonar calls and listen to their echoes arising from reflections of surrounding objects (Kössl et al., 2014; Moss and Surlykke, 2010; Simmons, 2012). Spectro-temporal parameters of echoes inform the animals about the position and identity of objects nearby (Wohlgemuth et al., 2016b). To gain spatial information, bats must assign incoming echoes to their corresponding calls (Corcoran and Moss, 2017; Suga et al., 1983; Ulanovsky et al., 2004). Call-echo assignments become challenging, however, when biosonar signals from many bats are overlapping (Corcoran and Moss, 2017; Levin et al., 2013; Parsons et al., 2003; Ulanovsky and Moss, 2008).

Under these circumstances, bats demonstrate a large repertoire of behavioral adaptations that are thought to represent behavioral strategies to improve signal extraction. These adaptations range from spectro-temporal changes in call design, to changes in call emission patterns (Adams et al., 2017; Amichai et al., 2015; Cvikel et al., 2015; Gillam and McCracken, 2007; Gillam et al., 2007; Habersetzer, 1981; Hage et al., 2013; Hiryu et al., 2010; Ibanez et al., 2004; Jarvis et al., 2013; Luo et al., 2015; Miller and Degn, 1981; Obrist, 1995; Ratcliffe et al., 2004; Roverud and Grinnell, 1985a; Roverud and Grinnell, 1985b; Simmons et al., 1979; Simmons et al., 1978; Takahashi et al., 2014; Tressler and Smotherman, 2009; Ulanovsky et al., 2004).

Our current understanding of why bats show such a large variety of adaptations when echolocating under “noisy” conditions is sparse. For example, it remains unknown whether the adaptations observed are individualistic and/or depend on the environmental context in which bats vocalize (i.e., the distance between the bat and the nearest target).

In this study, we tested the hypothesis that individual bats rely on different combinations of behavioral adaptations to overcome noise and that they can switch adaptation strategies at any timepoint during echolocation. To test this hypothesis and to gain a clearer understanding of echolocation behavior in “noisy” environments, individual bats of the species *Carollia perspicillata* were attached on the mass of a swinging pendulum (Figure 1A). The pendulum offers a behavioral paradigm whereby bats could actively echolocate in controlled scenarios, which could be replicated over several trials (Beetz et al., 2016b; Beetz et al., 2017; Henson et al., 1982; Macias et al., 2016). In our experiments, during forward swings – which mimicked a bat closing in on a target– the animals were acoustically stimulated with patterned echolocation calls broadcast from a speaker, which travelled with and pointing towards the animal (test trial). The call design and emission pattern of test trials were then compared to those recorded during control trials in which bats were swung in the absence of playback stimuli. During test trials, we examined whether bats would change different echolocation parameters, including call duration, call level, call frequency composition, and call emission pattern.

**Fig. 1.**
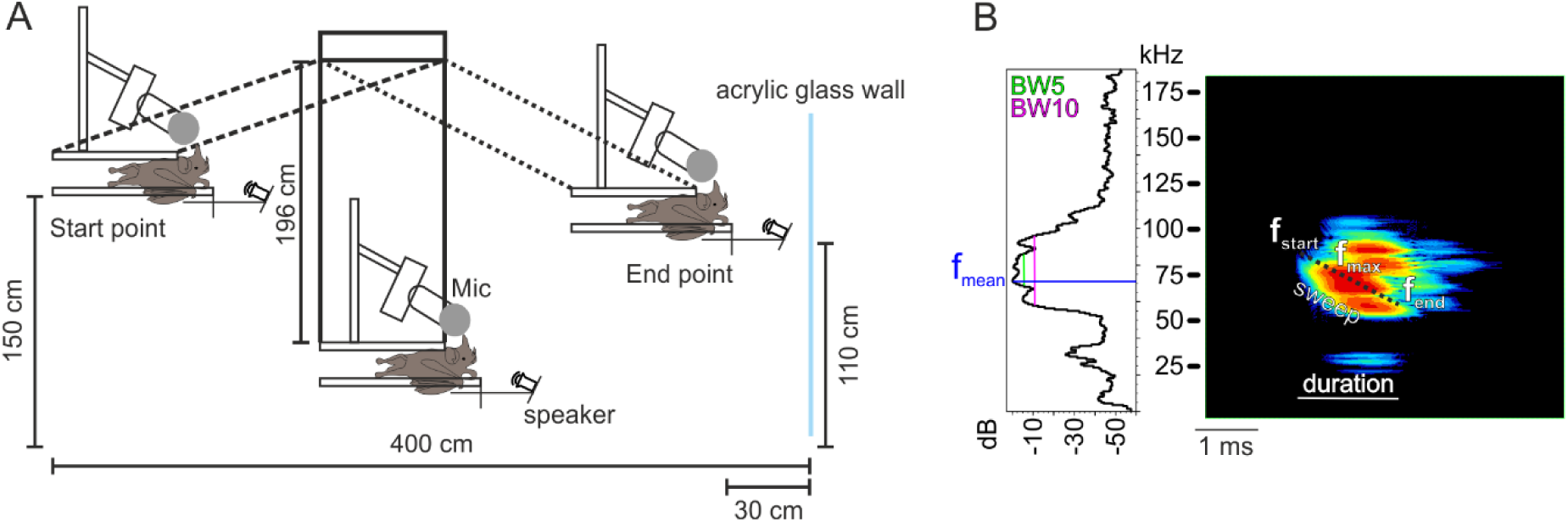
Behavioral paradigm and representative echolocation call. (A) Schematic side view of the pendulum paradigm. The bat was positioned on the mass of a pendulum, which was swung towards an acrylic glass wall. During the swing, the bat emitted echolocation calls that were recorded together with the echoes by an ultrasound microphone (Mic). For test trials, the bat was stimulated with playback echolocation sequences that were composed of a previously recorded echolocation call of the tested bat. The playback stimuli were emitted with a speaker that was pointing towards the bat’s head. Microphone and speaker were travelling with the bat and had a constant distance to the bat’s head throughout the experiments. (B) Power spectrum (left) and spectrogram (right) of a representative echolocation call recorded with the pendulum paradigm. Different call parameters were measured to characterize spectro-temporal call properties. Spectral parameters that were measured included initial (f_start_), centre (f_centre_), terminal (f_end_), mean (f_mean_), maximum amplitude (f_max_) peak frequency, and bandwidths at five (BW5) and ten dB (BW10) below the f_mean_. Call duration represents one of the temporal echolocation parameters that was considered in the analysis. The sweep rate represents the difference of f_end_ and f_start_ (f_end_ – f_start_) divided by the call duration.

## Materials and Methods

### Animals

Experiments were conducted on 10 bats (5 females and 5 males) of the species *Carollia perspicillata*. The bats were bred and kept in a colony at the Institute for Cell Biology and Neuroscience (Goethe-University Frankfurt). The experiments complied with all current German laws on animal experimentation and in accordance with the Declaration of Helsinki. All experimental protocols were approved by the Regierungspräsidium Darmstadt (experimental permit # #FU-1126).

### Pendulum paradigm and audio recordings

For controlling behavioral context, bats were positioned on the mass of a pendulum and were repetitively swung towards an acrylic glass wall (50 × 150 cm, Figure 1A) (Beetz et al., 2016b; Beetz et al., 2017; Henson et al., 1982; Macias et al., 2016). During the swing, the bats emitted echolocation sequences that were recorded, together with their echoes, by an ultrasound sensitive microphone (CM16/CMPA, Avisoft Bioacoustics, Germany). The microphone had a sensitivity of 50 mV/Pa and an input-referred self-noise level of 18 dB SPL, as reported by the manufacturer. The frequency response curve was flat (± 3 dB, as specified by the manufacturer) in the range from 30-130 kHz. The microphone travelled with the mass of the pendulum, which was medially positioned above the bat’s head. The membrane of the microphone was adjusted as closely as possible to the bat’s ears (∼ 4 cm). The microphone was connected to a sound acquisition system (Ultra Sound Gate 116Hm mobile recording interface, + Recorder Software, Avisoft Bioacoustics, Germany). To test the influence of acoustic interference on echolocation behavior, bats were swung in the pendulum while they were acoustically stimulated with a playback stimulus (see below). We compared the echolocation behavior recorded in the absence of playback stimuli (control trials) with the one shown in the presence of playback (test trials). Our reasoning was that because the behavioral context was invariant during control and test trials, except for the occurrence of the playback stimulus, we could correlate adaptations in the echolocation behavior with the presence/absence of the playback.

Initially, the bats were tested in a control trial followed by test trials where an echolocation call recorded during the forward swing of the control trial was selected to construct an individual-specific playback stimulus. The playback stimulus consisted of an echolocation call that was presented as quartets with a call interval of 25 ms and the quartets were repeated with an inter-quartet interval between 130 and 150 ms. The intensity of the playback stimulus was adjusted to rms values (of single calls) between 80 and 90 dB SPL for all animals. We reasoned that using an echolocation call of the tested animal, as a playback stimulus, would be the most effective way of achieving acoustic jamming. The latter is supported by the fact that subtle inter-individual differences in call design could be detected by the animals, which reduces signal interference (Yovel et al., 2009). During test trials, the playback stimulus was presented from an ultrasound speaker (MK 103.1 Microtech Gefell Microphone Capsule used as speaker) that was flat in the range from 5 to 120 kHz (mean level in calibration curve 84 ± 3 dB SPL, the speaker’s protection cap was replaced with a self-made cap to prevent energy loss at high frequencies). The speaker was placed pointing towards the bat’s head at a distance of 20 cm. The short distance between speaker and animal and the relatively tight fixation of the bat’s head prevented situations in which the bat could reduce acoustic interference via motor responses like head “waggling” (Wohlgemuth et al., 2016a). Thus, the bats had to rely mostly on changes in call design or emission pattern to minimize signal interference. Eight out of ten bats were tested on two consecutive days, but with different, day-specific, playback stimuli. The latter controlled for changes of the call design that may occur across days might bias our analysis. An overview of the call parameters used for constructing playback stimuli is shown in Table 1.

**Table 1.** Call parameters of the playback stimuli. Each animal was stimulated with one of its own echolocation calls to ensure a high probability of acoustic interference. Since some call parameters change across days within an individual, a new jamming stimulus was generated each day. BW = bandwidth; p f = peak frequency

| animal (day) | call duration [ms] | intensity rms [dB] | p f start [kHz] | p f end [kHz] | p f centre [kHz] | p f max [kHz] | p f mean [kHz] | BW5 [kHz] | BW10 [kHz] | sweep rate [kHz/ms] |
| --- | --- | --- | --- | --- | --- | --- | --- | --- | --- | --- |
| f8 (1) | 1,57 | 80,13 | 68,8 | 68,1 | 67,3 | 68,1 | 68,1 | 4,3 | 20,5 | -0,446 |
| f8 (2) | 1,62 | 83,77 | 82 | 82 | 82,7 | 82 | 82 | 9,5 | 12,4 | 0,000 |
| f9 (1) | 1,94 | 87,02 | 61,5 | 82 | 82 | 82 | 82,7 | 27 | 30 | 10,567 |
| f9 (2) | 1,64 | 78,8 | 80,5 | 90,8 | 87,1 | 87,1 | 87,1 | 11,7 | 27,8 | 6,280 |
| f10 (1) | 1,85 | 85,86 | 68,1 | 85,6 | 82,7 | 86,4 | 85,6 | 16,8 | 28,5 | 9,459 |
| f11 (1) | 2,34 | 84,9 | 74,7 | 76,9 | 82 | 82 | 82 | 21,2 | 29,2 | 0,940 |
| f11 (2) | 2 | 78,13 | 69,5 | 79,8 | 85,6 | 86,4 | 86,4 | 7,3 | 13,1 | 5,150 |
| f12 (1) | 1,96 | 85,6 | 74,7 | 79,1 | 79,1 | 85,6 | 79,1 | 19 | 24,1 | 2,245 |
| f12 (2) | 2,15 | 80,25 | 80,5 | 82 | 82,7 | 87,1 | 82,7 | 10,2 | 27 | 0,698 |
| m9 (1) | 2,04 | 87,08 | 79,1 | 84,2 | 82,7 | 71 | 71 | 33,6 | 38,8 | 2,500 |
| m9 (2) | 2,3 | 83,95 | 78,3 | 82,7 | 82,7 | 82,7 | 82,7 | 5,1 | 22,7 | 1,913 |
| m10 (1) | 2,21 | 86,72 | 71 | 80,5 | 80,5 | 81,2 | 80,5 | 5,8 | 10,2 | 4,299 |
| m10 (2) | 1,62 | 82,25 | 67,3 | 84,9 | 82 | 87,1 | 82,7 | 9,5 | 32,9 | 10,864 |
| m11 (1) | 1,68 | 81,53 | 70,3 | 81,2 | 82 | 82 | 82,7 | 5,1 | 10,9 | 6,488 |
| m12 (1) | 1,4 | 83,57 | 79,1 | 66,6 | 61,5 | 79,1 | 67,3 | 23,4 | 24,9 | -8,929 |
| m12 (2) | 1,53 | 77,23 | 79,8 | 89,3 | 87,8 | 87,8 | 87,8 | 4,3 | 27,8 | 6,209 |
| m13 (1) | 2,64 | 84,92 | 72,5 | 82 | 82,7 | 85,6 | 82,7 | 7,3 | 11,7 | 3,598 |
| m13 (2) | 2 | 76,35 | 67,3 | 81,2 | 80,5 | 82 | 80,5 | 10,9 | 33,6 | 6,950 |

### Analyzed echolocation parameters

Since the time pattern of the playback stimuli was kept constant, we could discriminate between biosonar signals emitted by the bat and the playback stimuli. The call emissions were manually tagged in the software Avisoft SAS Lab Pro (Avisoft Bioacoustics, Germany). To characterize the echolocation calls, different call parameters were measured in Avisoft SAS Lab Pro. The present study focused on call level, call duration, peak frequency at different call time points (start, end, maximum amplitude, and mean), bandwidth 5 (BW5), BW10, and sweep rate (Figure 1B). Regarding the call spectra, we considered the peak frequencies (frequencies with the maximum energy at particular time points of the call or on average of a call), because peak frequencies were likely to be the most salient spectral information of the echo that would suffer least from reflective attenuation. BW5 and BW10 represents frequency ranges at 5 and 10 dB below the mean peak frequency (Figure 1B). The sweep rate was calculated by subtracting the initial peak frequency from the terminal peak frequency and by dividing by the call duration.

The call emission pattern was characterized by measuring the call intervals and the tendency of grouping the calls. Analysis of the call groups was conducted using custom-written scripts in Matlab 2014 (MathWorks, USA). Call groups were defined according to two criteria (Beetz et al., 2018; Kothari et al., 2014). An “island criterion” defined call groups that were isolated in time. An isolation was fulfilled as soon as the preceding and following call intervals of a call group were 20% longer than the call intervals within call groups. If the “island criterion” was fulfilled, a second criterion, the so called “stability criterion”, defined the size of the call groups indicated by the number of calls belonging to a group. The stability criterion was fulfilled if the call intervals within call groups were stable with a 5% tolerance. Next, we calculated a strobe index for each animal and each condition (control and test trial). The strobe index represented the relative amount of calls that were emitted as groups.

### Statistics

For statistical analysis, we used the software GraphPad Prism 7 (GraphPad Software, USA; * p < 0.05; ** p < 0.005; *** p < 0.001; **** p < 0.0001). For analyzing distance-dependent changes of the echolocation behavior in the pendulum, non-parametric Kruskal-Wallis tests and a Dunn‘s multiple comparison post hoc tests were computed. For analyzing individual specific call adaptations in response to acoustic playback, control and test trials were directly compared from each animal by performing non-parametric Mann Whitney (in case of non-Gaussian distribution according to D’Agostino & Pearson normality test; alpha = 0.05) or parametric t-Tests (in case of Gaussian distribution according to D’Agostino & Pearson normality test; alpha = 0.05). For a comparison of the echolocation behavior between subsequent trials, non-parametric Kruskal-Wallis tests and Dunn‘s multiple comparison post hoc tests (in case of non-Gaussian distribution according to D’Agostino & Pearson normality test; alpha = 0.05) or ordinary one-way ANOVA and Tukey’s multiple comparison post hoc tests (in case of Gaussian distribution according to D’Agostino & Pearson normality test; alpha = 0.05) were computed.

## Results

### Pendulum paradigm mimics a natural approach flight

When swinging bats on the mass of a pendulum, they often emit echolocation calls (Beetz et al., 2016a; Henson et al., 1982; Macias et al., 2016). Thus, a pendulum paradigm allows to describe echolocation behavior under controlled conditions. This is important to test the influence of acoustic playbacks on echolocation behavior, independent from changes in the echolocation behavior due to target distances.

First, we quantified if pendulum-forward swings evoked consistent distance-dependent adjustments of the echolocation behavior in *C. perspicillata*. Based on 32 forward swings, each recorded from a different individual, we found that the bats shortened their call duration and inter-call intervals with decreasing target distance (Figure 2A-2B). In addition, with decreasing target distance, the bats increased their call intensity, starting peak frequency and peak frequency at the call’s maximum energy (peak freq max; Figure 2C-2E). Since the distance-dependent adjustments in call duration and call interval are comparable in the pendulum (laboratory condition) as in freely-flying bats (Thies et al., 1998), we concluded that a forward swing in the pendulum mimics a bat zooming in on a target in natural conditions.

**Fig. 2.**
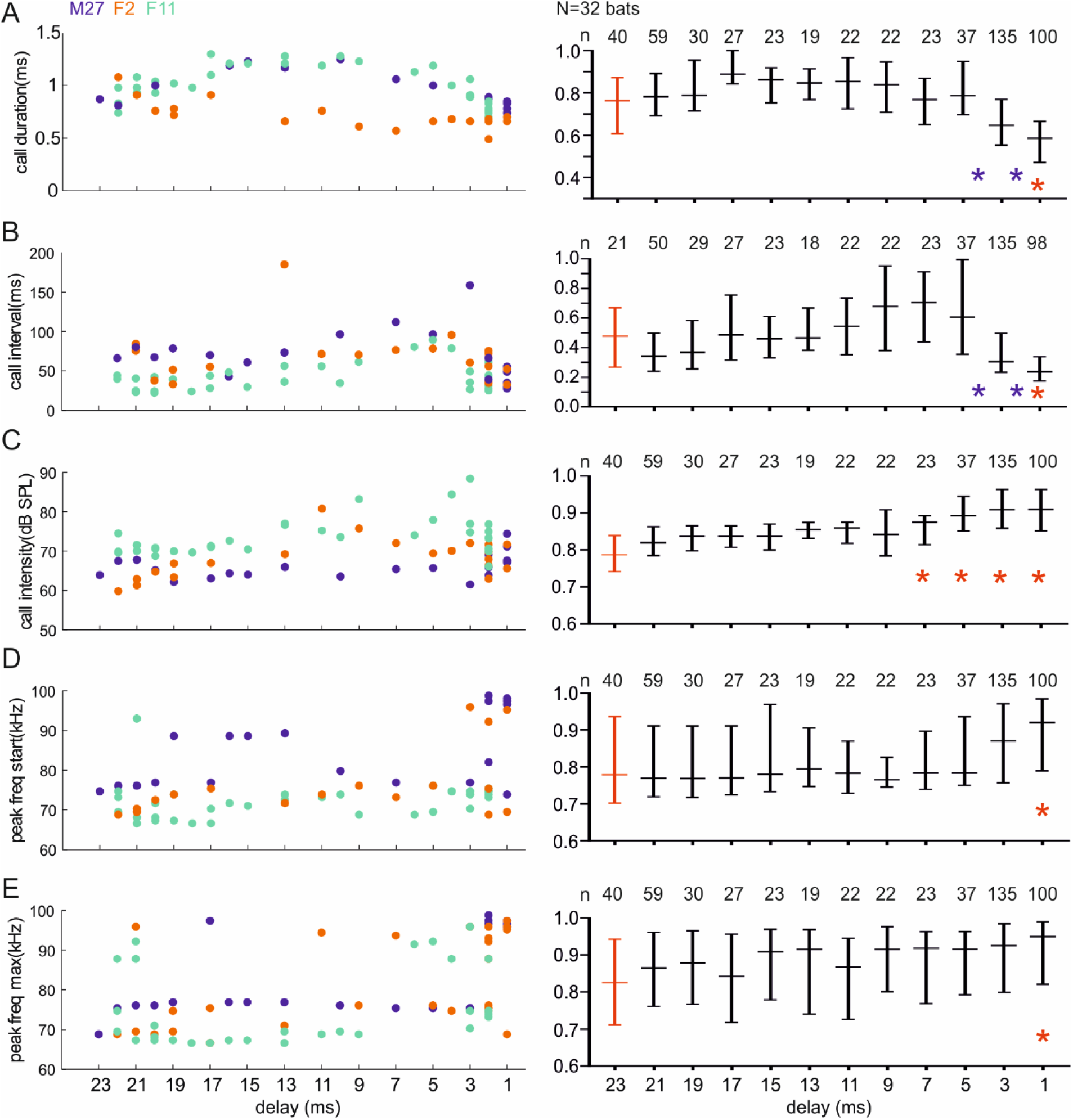
Distance-dependent changes of echolocation parameters during pendulum-forward swings. *C. perspicillata* reduces call durations (**A**) and call-intervals (**B**) with shorter distances to an object. Call intensities (**C**), initial peak frequency (**D**), and maximum peak frequency (**E**) slightly increase with shorter distance to an object. Subfigures on the left represent examples from three animals (M27, F2, F11) and subfigures on the right represent the median and the interquartile range from data of 32 bats in which each trial was normalized to its maximum value. Blue stars indicate significant differences (p < 0.05) between subsequent echolocation calls. Red stars indicate significant differences (p < 0.05) with calls emitted at echo delays between 23-22 ms (red data point). Kruskal-Wallis test + Dunn‘s multiple comparison post hoc test.

Next, we tested the inter-swing variability in the echolocation behavior of nine bats (five females, four males, Supplementary Table 1). Call duration, intensity, starting peak frequency, sweep rate, terminal peak frequency, peak frequency at call’s maximum energy, mean peak frequency, bandwidth 5, and bandwidth 10 did not vary over subsequent swings (Supplementary Table 1; p > 0.05) indicating that one forward swing reliably represents the echolocation behavior of an individual bat. Only four animals (female/F9, F10, F11, and male/M12) increased their call intervals across subsequent trials (Mann-Whitney test for F9 and F10; Kruskal-Wallis test for F11 and M12; p < 0.005), which may indicate that the bats habituated to the pendulum and therefore decreased the call rate.

### Individual bats change different call parameters in response to acoustic playback

To quantify adaptations of the echolocation behavior in response to acoustic playback, bats were swung in the pendulum while presenting an echolocation sequence (playback stimulus). The sequence was presented through a speaker attached to the pendulum mass, pointing towards the animal’s head (Figure 1A, test trials). One echolocation call from each tested bat served as building block for the playback stimulus (see methods for details). Thus, for each animal and experimental day, a new “individualized” playback stimulus was constructed (for stimulus details see methods and Table 1). In total, the echolocation behavior in the presence of playback stimuli was characterized in ten bats (5 females and 5 males). Echolocation behavior in the presence of playback was compared with the behavior recorded during an initial control trial in which no acoustic stimulus was played back to the animals. To minimize habituation to the pendulum paradigm, we decided to have only one control trial per session (per animal and day). As previously described, call parameters from subsequent control trials do not vary across swings (except for call intervals, Supple Table 1). Thus, one control trial is enough to characterize the bat’s echolocation behavior in the absence of playback stimuli. Since bats adjust their call design and emission pattern with the target distance (Figure 2), we pooled the calls into two groups, namely “long delay calls” and “short delay calls”. Echolocation calls that were broadcasted as the bat was farther than 1 m away from the acrylic glass wall were defined as “long delay calls”. Here, the echoes are delayed by more than 6 ms from the calls. Accordingly, echolocation calls that were emitted when the bat was closer than 1 m from the acrylic glass wall were defined as “short delay calls” (echo delays equal to or shorter than 6 ms).

In the presence of the playback stimulus, each individual bat demonstrated different combinations of adaptations (Table 2 and Supple Table 2). Four bats (F11, F12, M9, M12) increased the tendency of grouping their calls into call packs (exemplarily shown for F11 in Figure 3A; test trial, see also population data in Figure 3E). Six bats (F8, F10, F11, M10, M12, and M13) varied their call intervals. However, only a reduction of the call interval (observed in two bats, F11, M13) could be interpreted as an adaptation in response to the playback stimulus. Increased call intervals may be interpreted as habituation to the pendulum paradigm (see also Supple Table 1). Three bats (F8, M9, M13) increased and another three bats decreased (F9, M11, M12) call intensity during the test trials (Table 2; example in Figure 3B). Five bats changed their call duration, two shortened (F8, M11), two lengthened (M9, M10) and one shortened their “short delay calls” and lengthened their “long delay calls” (F9, Figure 3B; Table 2; Supple Table 2). The adaptation in call duration of F9 indicated that some bats differentially adapt “long delay calls” and “short delay calls” in response to playbacks. Changes in call spectra were sometimes prominent (Figure 3C and 3D) but also varied across animals (Table 2; Supplement Table 2). Calls shown in figure 3C and 3D were recorded as the bat had approximately the same distance to the target (∼2 m). Seven out of eight bats that changed their calls’ sweep rate decreased it when in the presence of playback. This indicated that the call frequency changed more slowly during the test compared to the control trials (Table 2; Supple Table 2). Changes in the sweep rate could be caused by changes of the call’s frequency range or by changes in call duration. Because lowering the sweep rate was not associated with lengthening the call, the sweep rate was mainly affected by changes in the frequency range. Seven animals (70%) changed either the BW5 or BW10 of the calls in the test trials. These changes could either be a BW decrease (shown by 40% of the bats tested; F8, F10, M11, M13) or an increase (shown by 30%; F9, F12, M9). Detailed data from three animals (A: F8; B: F9; C: M9) are plotted as boxplots in figure 4A-4C. For reasons of visualization, only call parameters that differed between the test and the control trials are plotted. Data from the remaining animals are presented in Supple figure 1. In conclusion, each animal adapted at least one call parameter in response to the playback stimuli. M11 was the only individual that did not change their call design (short delay calls) during the test trials. Overall, the bats changed different combinations of their call parameters, indicating that there was no common rule as to how to adapt to the playback stimuli.

**Table 2.**
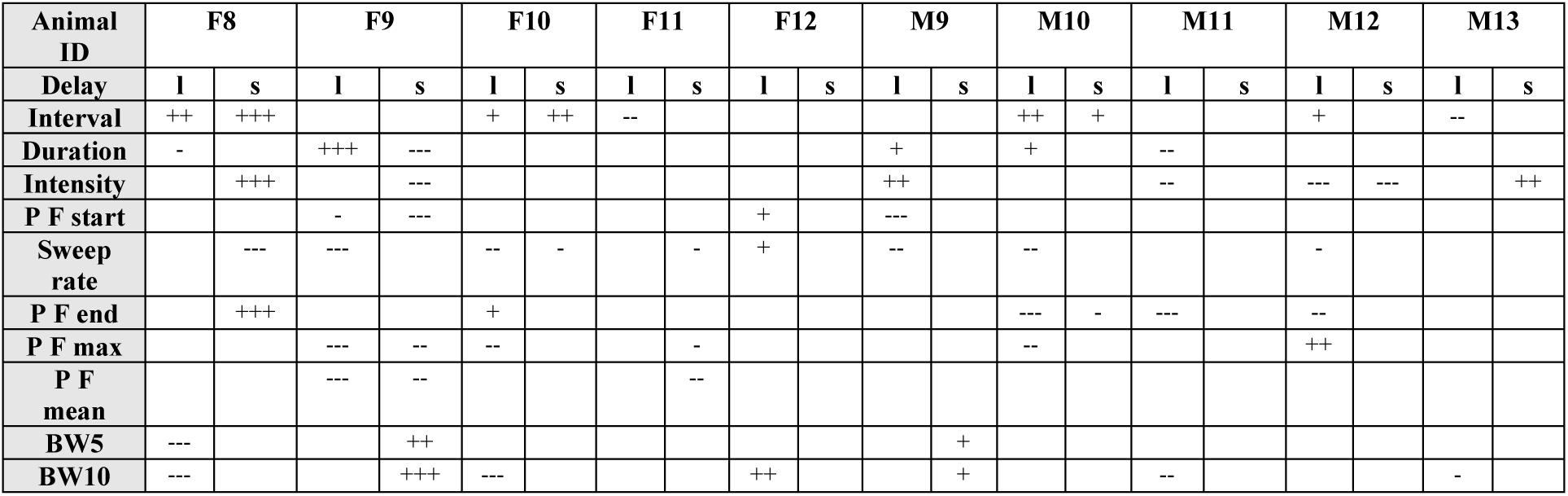
Changes of the call parameters induced by the presence of playback stimuli. + = higher values for test than for control trials (+ = p < 0.05; ++ p < 0.01; +++ p < 0.001); - = lower values for test than for control trials (- = p < 0.05; -- p < 0.01; --- p < 0.001); F = female; M = male; l = long delay calls; p f = peak frequency; s = short delay calls

**Fig. 3.**
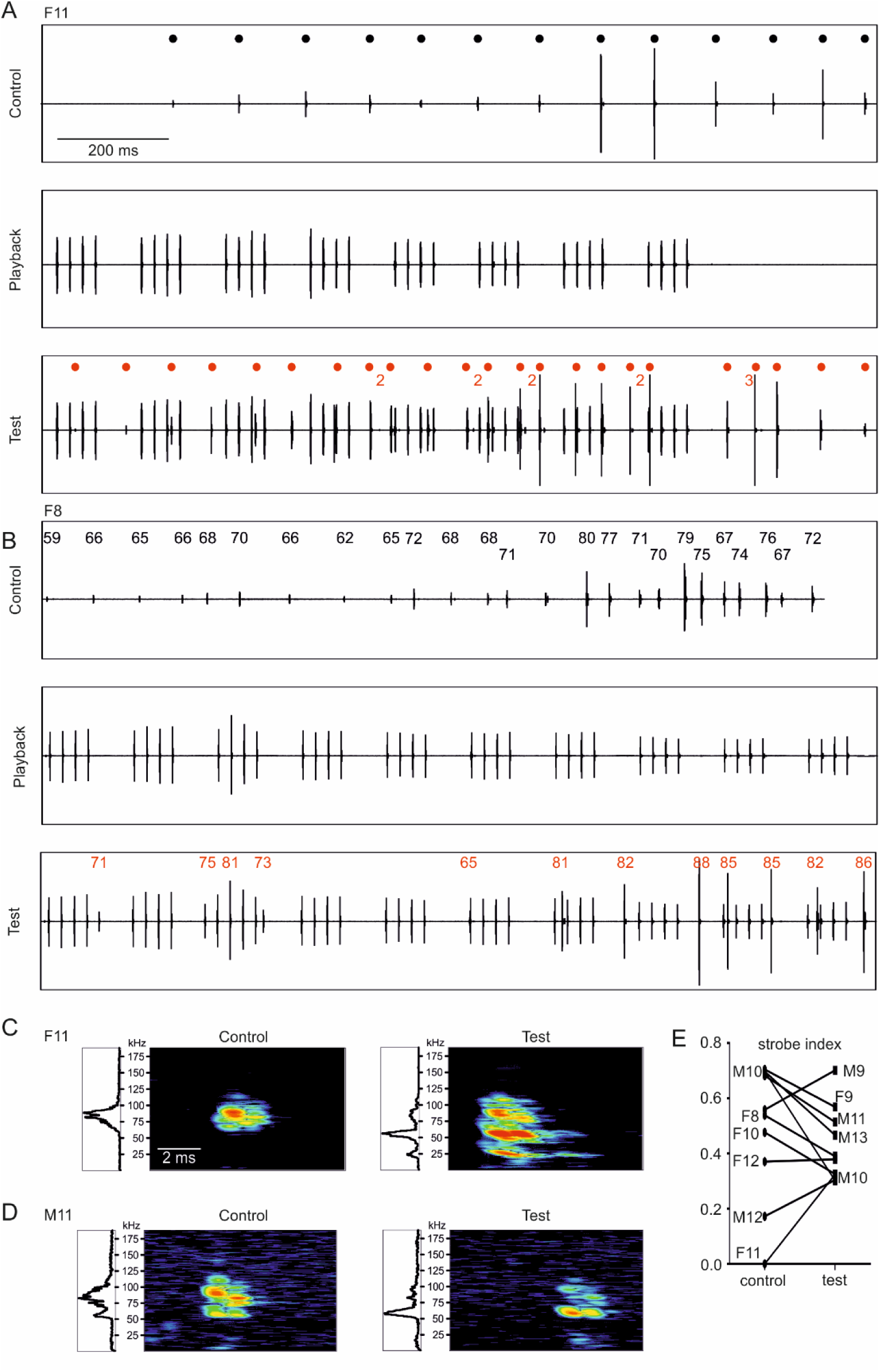
Examples of echolocation adaptation behaviors in response to playback stimuli. (**A**) Oscillograms of one control trial (top), the playback stimulus (middle), and one test trial (bottom) from female 11 (F11). Time points of call emissions are indicated by black or red dots above each oscillogram. During the control trial, the bat did not emit echolocation call groups. During the test trial, the bat grouped some calls into doublets (indicated by the number 2) or triplets (indicated by a “3”). Note that the jamming stimulus was recorded in addition to the echolocation calls of the test trial. Thus, oscillogram deflections without a dot represent signals coming from the playback stimulus. (**B**) Oscillogram of one control trial (top), the playback stimulus (middle), and one test trial (bottom) from female 8 (F8). In comparison to the calls emitted during the control trials, the call intensity was increased during the test trials. Numbers above each emitted call indicate the call intensity. (**C-D**) Power spectra (left) and spectrograms (right) of representative calls emitted during the control and test trial for two individuals (F11, M11). To exclude distance-dependent changes in the call design, all four calls were recorded as the bat was ∼2 meters away from the acrylic wall. Both bats decreased the bandwidth and mean peak frequency of their calls during the test trials as compared with the calls recorded during the control trials. (**E**) Tendency of emitting grouped calls (strobe index) under control and test conditions in all bats tested (n=10).

**Fig. 4.**
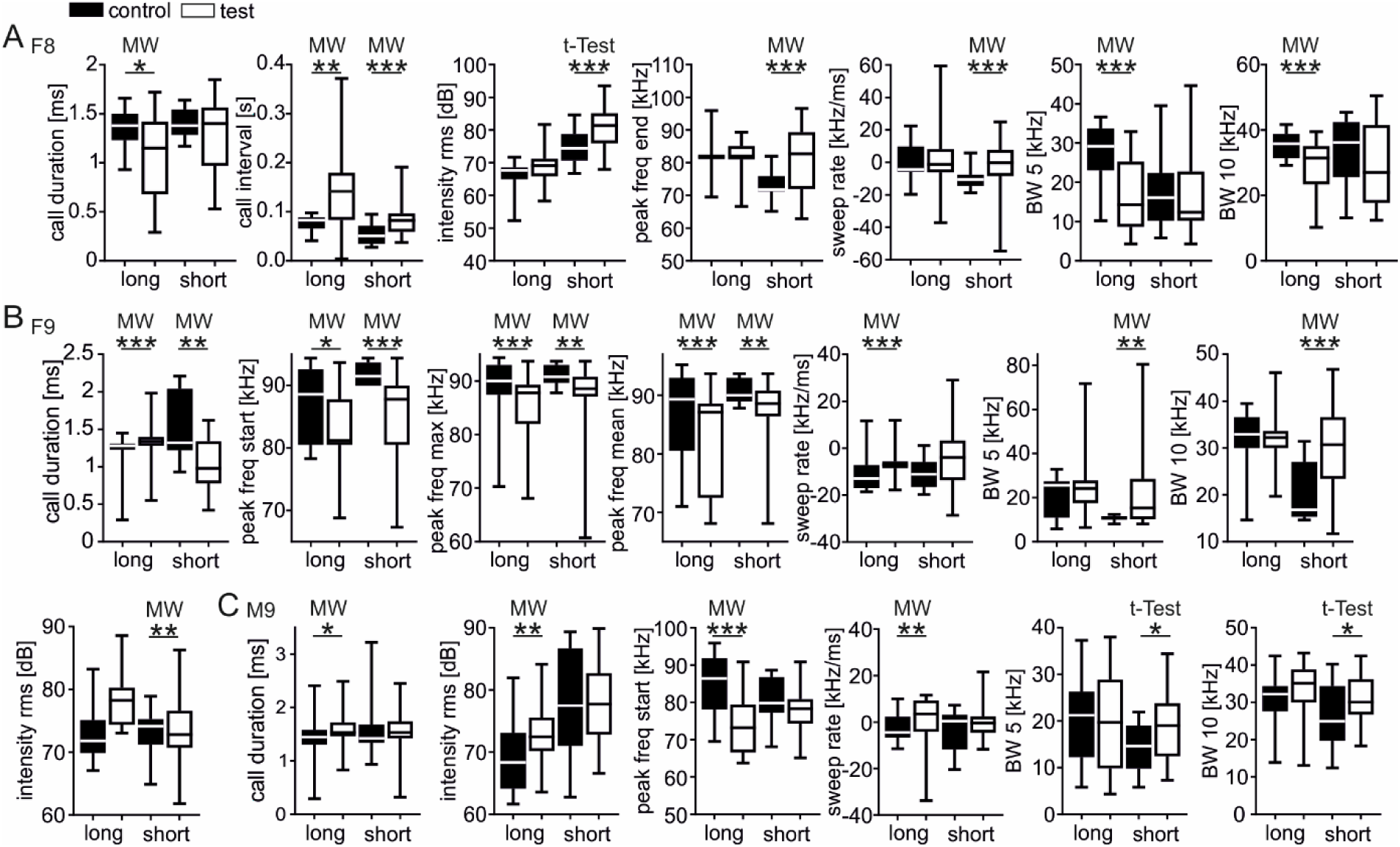
Individual specific call adaptations in response to playback stimuli. (**A-C**) Boxplots (whiskers represent minimum and maximum values) from three individuals (female 8 = 1_st_ row; female 9 = 2_nd_ row; male 9 = 3_rd_ row), showing call parameters that bats changed in response to the playback stimulus. Calls recorded under control conditions (absence of playback stimulus) are indicated by black boxplots, while white boxplots represent calls recorded under test conditions (presence of playback stimulus). Echolocation calls that are followed by an echo within 6 ms were grouped into “short delay calls”. Echoes following a call by more than 6 ms were grouped into “long delay calls”. Note that each bat changed their different call parameters under test conditions. MW = Mann-Whitney test; * p < 0.05; ** p < 0.005; *** p < 0.001.

### Bats vary adaptation strategies across trials and days

We were interested in assessing if each individual bat prefers the same combination of adaptations or whether the bats change their strategies across days or even across trials on the same day. However, before characterizing the temporal dynamics of the adaptations, we quantified the variability of the call design across subsequent days under controlled conditions (absence of playback stimulus). We tested ten bats in the absence of playback stimuli for two (F1, M3, M7) or three (F2, F3, F5, F7, M1, M4, M6) subsequent days (Supple Table 3). Although the bats did not vary their call design across subsequent control trials on the same day (Supple Table 1), they dramatically varied their call design across subsequent days (Supple Table 3). Thus, to test if bats change their adaptations in response to the playback stimulus across days, we recorded an initial control trial on each day and compared the echolocation behavior from the day-specific control trial with the one recorded during test trials.

**Table 3.** Changes of the call parameters across trials. + = higher values for test than for control trials (+ = p < 0.05; ++ p < 0.01; +++ p < 0.001); - = lower values for test than for control trials (- = p < 0.05; -- p < 0.01; --- p < 0.001; p < 0.0001); F = female; M = male; l = long delay calls; p f = peak frequency; s = short delay calls

| Animal ID<br>(calls/trial) | Trial<br>(Day) | Intensity | Duration | Interval | Sweep<br>rate | P F<br>start | P F<br>end | P F<br>max | P F<br>mean | BW5 | BW10 |
| --- | --- | --- | --- | --- | --- | --- | --- | --- | --- | --- | --- |
| F8 (25, 16, 14, 12,<br>16, 16, 13, 11) | 1 (1st) | + | --- | +++ | - |  | ++ |  |  | -- | --- |
|  | 2 (1st) |  |  |  |  |  |  | + |  |  |  |
|  | 3 (1st) | ++ |  | ++ |  |  |  |  |  | -- | ---- |
|  | 4 (2nd) |  |  |  | - |  | + |  |  |  |  |
|  | 5 (2nd) |  |  |  |  |  |  |  |  |  |  |
|  | 6 (2nd) |  |  | + |  |  |  |  |  |  |  |
| F9 (28, 15, 10, 15,<br>17, 23, 18, 34, 16,<br>16) | 1 (1st) |  |  |  |  | - |  |  |  |  |  |
|  | 2 (1st) | - |  |  | - | -- |  | -- |  |  |  |
|  | 3 (1st) |  |  |  |  | - |  | -- | - |  |  |
|  | 4 (1st) |  | +++ |  |  | -- |  | ---- | --- | + | + |
|  | 5 (1st) |  | + |  |  | -- |  | - | - |  |  |
|  | 6 (2nd) | ++ |  |  | -- |  | +++ |  |  |  |  |
|  | 7 (2nd) |  |  | + |  |  |  |  |  |  |  |
|  | 8 (2nd) | - |  | + | - |  | ++ |  |  |  |  |
| F10 (22, 13, 10,<br>22, 16, 15, 16) | 1 (1st) |  |  |  | -- |  |  |  |  |  |  |
|  | 2 (1st) |  | + |  | -- |  |  | -- |  |  |  |
|  | 3 (1st) |  |  |  | - |  |  |  |  |  |  |
|  | 4 (1st) |  |  |  | - |  |  | - |  |  |  |
|  | 5 (1st) |  |  |  |  |  | + |  |  |  |  |
|  | 6 (1st) |  |  | + |  |  |  |  |  |  |  |
| F11 (13, 23, 7, 20,<br>8, 9, 16, 13, 11,<br>14, 15, 10, 9, 11) | 1 (1st) | + |  | --- |  |  |  | - |  |  |  |
|  | 2 (1st) |  | - |  |  |  |  |  |  |  |  |
|  | 3 (1st) |  |  | - |  |  | - | -- | -- |  |  |
|  | 4 (1st) |  |  | - |  |  |  | - | - |  |  |
|  | 5 (2nd) |  |  | - |  |  |  |  |  | ++ |  |
|  | 6 (2nd) |  |  |  |  |  |  |  |  |  |  |
|  | 7 (2nd) |  |  |  |  |  |  |  |  |  |  |
|  | 8 (2nd) |  |  |  |  |  |  |  |  |  |  |
|  | 9 (2nd) |  |  |  |  |  |  |  |  |  |  |
|  | 10 (2nd) | -- |  |  |  |  |  |  |  |  |  |
|  | 11 (2nd) |  |  | -- |  |  |  |  |  |  |  |
|  | 12 (2nd) |  |  |  |  |  |  |  |  |  |  |
| F12 (18, 14, 11,<br>11, 11, 24, 17, 16) | 1 (1st) |  | + |  |  |  |  |  |  |  |  |
|  | 2 (1st) |  |  | ++ | - |  |  |  |  |  | + |
|  | 3 (1st) |  |  |  |  |  | - |  |  |  |  |
|  | 4 (2nd) |  |  | ---- | ++++ | ++ | -- |  |  | ++ | ++ |
|  | 5 (2nd) |  |  | - | + |  | - |  |  | + | + |
|  | 6 (2nd) |  |  | - | + |  | - |  |  | + | ++ |
| M9 (23, 24, 15, 7,<br>9, 24, 23, 9, 7) | 1 (1st) | ++ | ++ |  | - | ---- |  |  |  |  |  |
|  | 2 (1st) |  |  | + |  | - |  |  |  |  |  |
|  | 3 (1st) | ++++ |  |  |  |  |  |  |  | --- |  |
|  | 4 (1st) |  |  |  | -- | ---- |  |  |  |  |  |
|  | 5 (2nd) |  |  | -- | - | -- |  |  |  | + | ++ |
|  | 6 (2nd) |  |  |  |  |  |  |  |  | + | + |
|  | 7 (2nd) |  |  |  |  |  |  |  |  | + |  |
| M10 (17, 7, 14,<br>18, 12, 12, 7, 7) | 1 (1st) |  |  |  |  |  | - | -- | - |  |  |
|  | 2 (1st) |  |  |  |  | -- |  | ---- |  | - |  |
|  | 3 (2nd) |  |  |  |  |  | - |  |  |  |  |
|  | 4 (2nd) | +++ | +++ | + |  |  |  |  |  |  |  |
|  | 5 (2nd) |  | + | ++ |  |  |  |  |  |  |  |
|  | 6 (2nd) |  | ++ | + | ++ | ++ | ---- |  |  |  |  |
| M11 (22, 8, 12,<br>11, 10) | 1 (1st) | -- |  |  |  |  | --- |  | -- |  |  |
|  | 2 (1st) |  |  |  |  |  | - |  |  |  |  |
|  | 3 (1st) |  |  |  |  |  |  |  | - |  |  |
|  | 4 (1st) | --- | - |  |  |  |  |  |  |  |  |
| M12 (20, 10, 7,<br>11, 13, 11) | 1 (1st) |  |  |  |  |  | - |  |  |  |  |
|  | 2 (1st) |  |  | ++ |  |  |  |  |  | -- | - |
|  | 3 (2nd) |  |  |  |  |  |  |  |  |  |  |
|  | 4 (2nd) |  |  |  | ++ |  |  |  |  |  |  |
| M13 (15, 29, 11,<br>24, 20, 20, 14, 16,<br>12, 15) | 1 (1st) | ++ |  | ---- | --- | +++ |  |  |  |  |  |
|  | 2 (1st) |  |  |  |  |  |  |  |  |  |  |
|  | 3 (1st) |  |  | - | --- | ++++ |  |  |  |  |  |
|  | 4 (2nd) |  |  |  | --- | -- |  |  |  | -- |  |
|  | 5 (2nd) |  |  | - | --- | -- |  |  |  |  |  |
|  | 6 (2nd) |  |  |  |  |  |  |  |  |  | - |
|  | 7 (2nd) |  | ---- | + |  |  | +++ |  |  |  |  |
|  | 8 (2nd) |  |  |  | -- | - | + |  |  | - |  |

Moreover, to perform a trial-by-trial analysis and to gather enough data points for statistical analysis, we pooled data from long and short delay calls. During the test trials, bats emitted slightly fewer calls than during control trials (median n of calls: 16.5 control and 13 test; Mann-Whitney test: p = 0.036). By comparing the call parameters from F9 across days (Figure 5; Table 3), it became clear that the adjustments of call duration (Figure 5C), starting (Figure 5B), maximum (Figure 5F), and mean peak frequency (Figure 5H), bandwidth 5 (Figure 5G) and bandwidth 10 (Figure 5I) exclusively occurred on day 1. On day 2, bat F9 mainly changed call intensity (Figure 5A), terminal peak frequency (Figure 5D), and sweep rate (Figure 5J). As already mentioned, increments in call interval (Figure 5E) did not necessarily represent an adaptation to reduce acoustic interference; possibly, they represented habituation to the pendulum paradigm across trials/days.

**Fig. 5.**
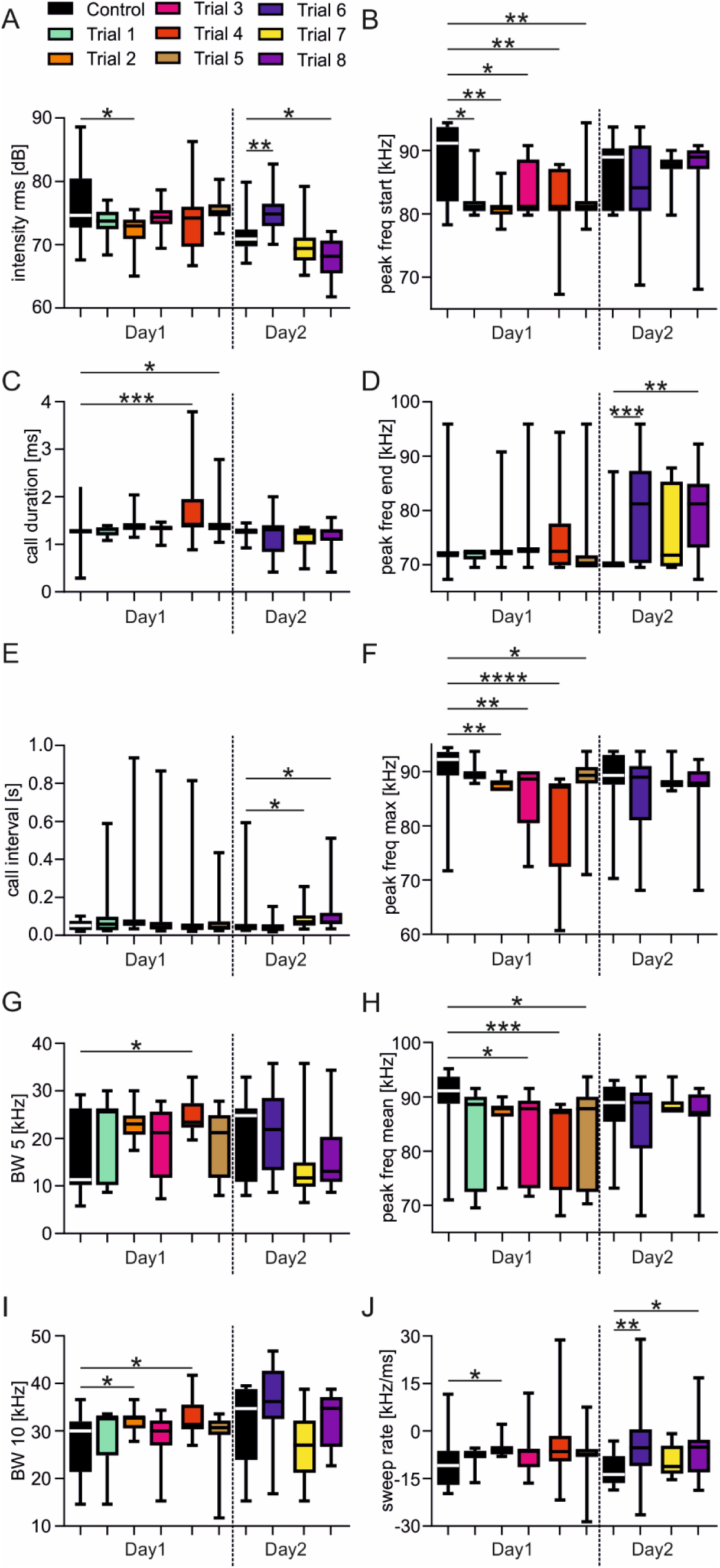
Bats switch adaptation strategies across trials and days. Call parameters are shown as boxplots (whiskers represent minimum and maximum values) for each trial (8 test trials and 2 control trials) across two days (from one bat). For visualization purposes, each trial is color coded and the control trials are shown in black. Note that the bat changes some call parameters only at day 1 (e.g., peak freq start; call duration; peak freq max) and not at day 2. Kruskal-Wallis Test and Dunn’s multiple comparison Post hoc test; * p < 0.05; ** p < 0.005; *** p < 0.001, **** p < 0.0001.

We observed that echolocation adaptation strategies not only varied across days, but also across subsequent trials (Table 3). For example, F9 changed the calls’ mean peak frequency in three (trial 3, 4, 5) out of five trials at day 1 (Figure 5H). Changes of other call parameters varied less dramatically across trials of the same day. In all trials on day 1, F9 decreased its starting (Figure 5B) and maximum peak frequency (Figure 5F). For detailed data from the remaining nine animals see S2-S10 figure. Overall, we found in 56 out of 67 test trials (83.6%) statistically significant differences between the control and test trials (Table 3). In eleven test trials, the bats did not change any call parameter compared to the control trial.

### Bats dynamically switch adaptation strategies within trials

What could have happened during test trials when we could not find an adaptation in the echolocation behavior? For these trials, was the acoustic interference too weak to evoke adaptations? Alternatively, might the bats have dynamically changed their adaptations during trials, so that the adaptation would not be detectable when pooling calls from an entire swing? To assess the latter idea, we compared parameters of each call from the test trial with the same parameters in the call used to construct the playback stimulus. The upper color maps, in figure 6A and 6B, exemplarily show the relative differences between call parameters and the playback parameters for two trials in two different bats (M9 and F12). The calls are ordered along columns in which the leftmost column represents the call with the longest echo-delay and the rightmost column represents the call with the shortest echo-delay. Each line represents the relative difference of a call parameter with respect to a playback parameter. This result was calculated by subtracting the playback parameters from the call parameters and by normalizing the difference against its absolute maximal difference for the entire trial. The darker the red and blue patches are, the more positive and negative were the call parameters in comparison to the playback stimulus. Based on the trial in figure 6A, the bat initially emitted calls with lower starting peak frequencies (peak start) and call intensities than the playback stimulus. At an echo delay of ∼3 ms (between the 12^th^ and 13^th^ call, white dashed line in Fig. 6A), the bat abruptly switched the strategy and increased the maximum and mean peak frequency while decreasing the BW of subsequent calls.

**Fig. 6.**
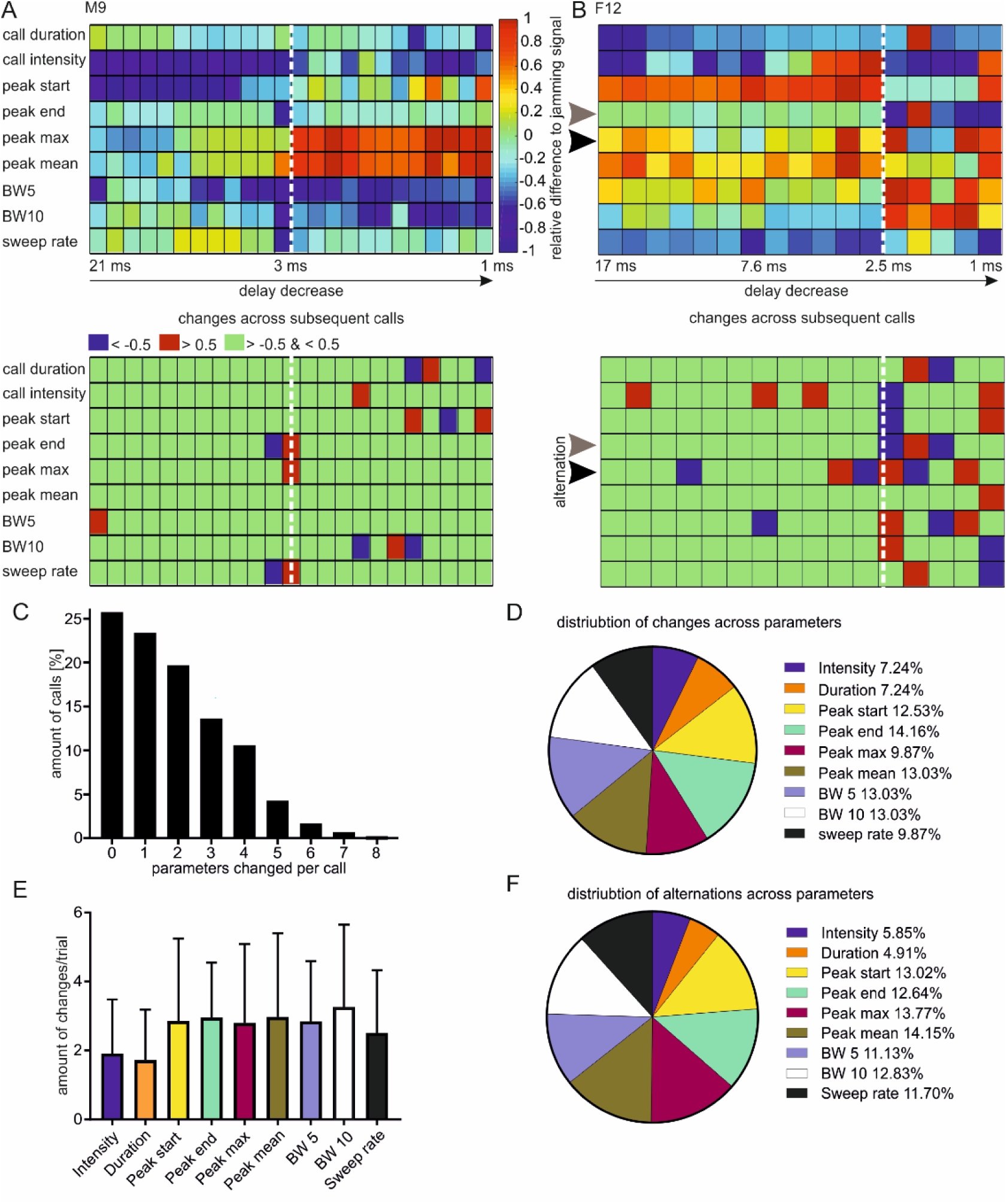
Bats dynamically change echolocation parameters within trials. (**A-B** upper graphs) Color maps from two representative test trials (M9 in (A) and F12 in (B)), illustrating the differences between calls and playback stimuli in a call-wise manner. Along the x-axis, the calls are ordered according to their emission order during the trial. The echo delay value from some call-echo pairs are indicated in the x-axis. Along the y-axis, normalized call parameter differences are color coded. The differences were normalized to their absolute maximum value at the corresponding parameter for the specific trial. The differences of the following call parameters were considered: call duration, call intensity, peak frequency at the beginning, end and maximum of the call, mean peak frequency of the call, bandwidth 5 (BW5), bandwidth 10 (BW10), and sweep rate. In some trials, a clear transition of the adaptation strategies can be detected (white vertical dashed lines). In some cases, the bats alternate call values, as exemplified for F12 for the terminal and maximum peak frequency indicated by a gray and black arrowhead, respectively. (**A-B** lower graphs) Colormaps illustrating abrupt changes of call parameters across subsequent calls. Abrupt changes occurred when a call parameter between two consecutive calls varied by more than 50% (blue and red cells represent reductions or increases in the corresponding call parameter). Changes of the call parameters that are below 50% were not abrupt enough to be defined as a change (green cells). Transitions between adaptation strategies and alternations between call parameter values can be seen more easily in the lower colormaps. (**C**) Histogram showing the level of parameters that are abruptly changed per call for all investigated calls (n = 889). Note that almost 75% of the calls show at least one abrupt change. (**D**) Pie chart illustrating the distribution of abrupt changes over the call parameters. Abrupt changes occur primarily within the call spectrum and less often for the intensity or duration. (**E**) Mean values of the amount of change per trial, plotted against the call parameter. Spectral parameters are shown to vary more often across trials than non-spectral ones (duration and intensity). (**F**) Pie chart representing the relative distribution of alternations across different call parameters.

To visualize abrupt changes better, we calculated the differences of the parameters of subsequent calls and plotted the values in the bottom color maps shown in figure 6A and 6B. We defined an abrupt change when the considered parameter varied by more than 50% between subsequent calls. For example, according to figure 6A, the terminal (peak end), maximum peak frequency (peak max), and sweep rate of call 13, are more than 50% higher than the ones of call 12. This outcome is indicated by red cells at the corresponding column (white dashed line) in the lower color map of figure 6A.

Abrupt changes in call design were also visible in other trials, like the one presented in figure 6B. Here, abrupt changes mainly occurred at around 2.5-ms echo delay (white dashed line) by decreasing the call intensity, starting (peak start), and terminal frequency (peak end) while the maximum peak frequency (peak max) as well as the call bandwidths (BW5 and BW10) abruptly increased. By considering all calls (889 calls from 69 trials and 10 animals), about three quarters of the calls (74.24%) showed abrupt changes in at least one call parameter (Figure 6C). About half of the calls (50.84%) showed abrupt changes in more than one call parameter. The bats did not focus on a specific call parameter, but rather changed most of their parameters with equal probability (Figure 6D). Call intensity and call duration were least (7.24%) abruptly changed within the trials.

When taking a closer look on the pattern of call changes over subsequent calls (color maps at the bottom of figure 6B), it became clear that the bats sometimes changed the call parameters in an alternating manner. During the second half of the trial, the bat alternated between high and low terminal (peak end) and maximum peak frequencies (peak max), indicated by gray and black arrowheads, respectively. Before analyzing the alternations in more detail, we questioned how often the bats changed a certain call parameter during the trial. The bar plot in figure 6E shows that the bats changed spectral parameters more often per trial (mean of peak start = 2.85 ± 2.39; mean of peak end = 2.96 ± 1.59; mean of peak max = 2.8 ± 2.29; mean of BW5 = 2.84 ± 1.75; mean of BW10 = 3.26 ± 2.39; mean of sweep rate = 2.51 ± 1.82) than the call intensity (mean = 1.91 ± 1.57) and the call duration (mean = 1.73 ± 1.46) (p < 10^−5^ Kruskal-Wallis test). Because spectral parameters varied more often during the trials, alternations occurred with a higher probability in spectral than in non-spectral (call intensity and call duration) parameters (Figure 6F). Across the spectral parameters, the probability of alternations did not differ significantly (p = 0.91 Kruskal-Wallis test), indicating that alternations could equally occur in each of the analyzed call parameters.

## Discussion

The present study characterized adaptation strategies of echolocating bats (fruit-eating bat *C. perspicillata*) in the presence of playback stimuli. These playback stimuli potentially interfered with the bat’s biosonar signals, making signal extraction for the bat challenging. In their natural environments, adjustments to echolocation behavior not only occur in the presence of an acoustic interferer, but also when bats approach obstacles or transition between different locales. Thus, it was crucial to test for the influence of acoustic interference under an otherwise invariant behavioral context. The pendulum paradigm, indeed, fulfilled these requirements because the behavioral scenario of an approach flight could be repetitively mimicked (Figure 2).

Our results demonstrate that *C. perspicillata* changes its call parameters and emission pattern when echolocating in the presence of playback stimuli. Interestingly, instead of relying on one adaptation strategy, the bats combined different adaptation strategies (Table 2 and Figure 4). To our surprise, the bats switched between different strategies across (Table 3, Figure 5) and even within trials (Figure 6). This flexibility renders the echolocation behavior, in the presence of acoustic interferers, highly dynamic and unique across different individuals and time points. Utilizing such dynamics, the bats can create unique echolocation streams that may be distinguished from interfering signals.

### Coping with signal interference

Signal interference is a challenge with which every animal must cope; they must extract behaviorally relevant signals from a noisy background. The greater the similarity between relevant signals and background, the more challenging signal extraction becomes. To facilitate signal extraction, animals show large repertoires of different behavioral adaptations (Corcoran and Moss, 2017; Ulanovsky and Moss, 2008) like orienting their sensory organs towards relevant signals (Eckmeier et al., 2008; Ganguly and Kleinfeld, 2004; Land, 2015; Ribak et al., 2009; Schroeder et al., 2010; Tarsitano and Andrew, 1999; Towal and Hartmann, 2006). For example, bats increase head waggles and the inter-pinna distance when orienting under challenging conditions (Wohlgemuth et al., 2016a). Potentially, this response improves localization of the echo source (Wohlgemuth et al., 2016a). Additionally, adjustments of the pinna’s shape and orientation may increase the directionality of hearing (Gao et al., 2011). In the present study, head waggles were limited by tightly positioning the bats on the platform of the pendulum mass. Moreover, by placing the jamming source close to the animals’ head, motor responses would barely facilitate signal extraction under such conditions.

For some behaviors - like communication, electrolocation, or echolocation - animals produce behaviorally relevant signals, which allow them to directly control the signal’s discriminability from its background. The latter becomes clear when considering the cocktail party problem (Bee and Micheyl, 2008). In a noisy environment, we can focus on our communication partner by carefully listening to him/her and improve the signal-to-noise ratio by increasing our voice intensity (Brumm and Zollinger, 2011; Luo et al., 2015), an adaptation known as the Lombard-effect. Signal extraction may not only be improved by increasing the signal-to-noise ratio but also by limiting the spectral overlap between signal and background. This adaptation has originally been described in electrolocating fish (Bullock et al., 1972; Watanabe and Takeda, 1963). When encountering animals whose signal frequencies overlap with the fish’s own signal frequency, the animals shift the signal frequencies away from each other. This behavior has been circumscribed as the Jamming Avoidance Response (JAR) and it reduces the signal interference with signals coming from conspecifics. JAR has also been demonstrated in different bat species (Gillam and McCracken, 2007; Gillam et al., 2007; Habersetzer, 1981; Hage et al., 2013; Ibanez et al., 2004; Miller and Degn, 1981; Ratcliffe et al., 2004; Takahashi et al., 2014; Tressler and Smotherman, 2009; Ulanovsky et al., 2004) and the present study. However, in contrast to weakly electric fish, which try to occupy an individual specific frequency band, bats dynamically adjust their call spectra in various situations. Bats adjust their calls when approaching an obstacle or when transiting between different habitats (Barchi et al., 2013; Falk et al., 2014; Griffin, 1953; Hiryu et al., 2010; Kalko, 1995; Kalko and Schnitzler, 1989; Knowles et al., 2015; Kothari et al., 2014; Petrites et al., 2009; Roverud and Grinnell, 1985a; Schnitzler et al., 1987; Simmons et al., 1978; Surlykke and Moss, 2000; Wheeler et al., 2016). Since frequency adjustments occur frequently and under various conditions, an adaptation that purely depends on a JAR may not be sufficient to orient collision-free in the presence of signal interferers. This hypothesis gains support by recent simulations that have tested for the effectiveness frequency adjustments when navigating in noisy environments (Mazar and Yovel, 2019). Note that some studies reported that bats do not shift their frequency in response to acoustic interference (Götze et al., 2016) or that the frequency shifts are correlated with the object’s distance rather with the presence of an acoustic interferer (Cvikel et al., 2015). Because we compared echolocation calls that were emitted roughly at similar object-distances, we can exclude that frequency shifts, present in our study, reflect distance-dependent changes of the call design.

### Repertoire of behavioral adaptations in response to interfering signals and their possible neural correlates

In the present study, most biosonar adjustments occurred in the spectral domain, although others were also detected in the temporal or energy domain (Figure 5D). Each adaptation may facilitate signal extraction in noisy environments. For example, decreasing the call bandwidth may reduce the spectral overlap between the playback stimulus and the biosonar signals. An increase in call bandwidth may recruit neurons that are not sensitive to the playback stimulus and therefore “selectively” process frequencies that are not occupied by the interferers. Bats also increase the signal-to-noise ratio by increasing call intensity (Amichai et al., 2015; Hage et al., 2013; Luo et al., 2015; Simmons, 2017; Simmons et al., 1978; Takahashi et al., 2014; Tressler and Smotherman, 2009). Unexpectedly, in the present study, sometimes the bats decreased their call intensity when they echolocated in the presence of interfering signals. Although this decreases the signal-to-noise ratio, it could still be useful from the perspective of neuronal processing. Many auditory neurons are more sensitive to low rather than to high sound levels (Barone et al., 1996; Hechavarría and Kössl, 2014; Park and Pollak, 1993; Suga and Manabe, 1982; Yang et al., 1992). This attribute makes the neurons highly selective to faint biosonar signals while being insensitive to intense background stimuli. Some studies have reported that bats lengthen their calls when flying in noisy environments (Amichai et al., 2015; Simmons, 2017; Simmons et al., 1979; Simmons et al., 1975; Tressler and Smotherman, 2009). In the present study, we observed that some bats lengthened, and others shortened their calls. Both adaptations putatively minimize acoustic interference. Shortening the calls decreases the chance of a temporal overlap between biosonar signals and the background. Lengthening the calls increases the risk of temporal overlap, but it could still be useful if only a small portion of the echo needed to be detected to gain enough spatial information.

Furthermore, not only the call design, but also the emission pattern can be adjusted to reduce or even avoid signal interference. Some bat species alternate between two call designs that differ in their frequency spectrum (Obrist, 1995; Roverud and Grinnell, 1985a; Roverud and Grinnell, 1985b). This adaptation allows a higher call rate by emitting a pair of calls before receiving an echo from the first call of the pair (Behr and von Helversen, 2004; Jung et al., 2007). The arising echoes differ in their frequency spectra which makes their discrimination feasible (Hiryu et al., 2010). Alternation of spectral call parameters have also been observed in the present study (Figure 5B, 5F). However, these alternations occurred occasionally and not throughout the entire trial. Thus, the behavioral importance of alternating call parameters in *C. perspicillata* needs to be further assessed.

Some bats reduce their call rate (Adams et al., 2017) and temporally even cease to emit calls (Jarvis et al., 2013). This adaptation may be beneficial if the bats eavesdrop on echolocation signals from conspecifics and use the signals for orientation (Barclay, 1982; Chiu et al., 2008; Leonard and Fenton, 1984; Lin and Abaid, 2015). Although, *C. perspicillata* emitted fewer calls during test compared to control trials, the pendulum paradigm was not designed to test for eavesdropping on the playback stimulus.

Lastly, some bats increase their rate of grouping calls when orienting in cluttered or noisy environments ((Beetz et al., 2018; Beetz et al., 2019; Luo et al., 2015; Roverud and Grinnell, 1985a) and present study). Grouping calls may improve echolocation performance in different ways. First, a defined periodicity of echo arrivals allows echo identification based on prediction (Petrites et al., 2009; Suga et al., 1983; Wheeler et al., 2016; Wohlgemuth et al., 2016a). Second, grouping the calls could create an information redundancy allowing the bats to rely only on a small portion of the call group (Beetz et al., 2018).

### Bats show different combinations of adaptations when echolocating in the presence of an acoustic interferer

Instead of adjusting just one echolocation parameter, when echolocating in noisy environments, our results indicate that bats have a toolbox of different and combinable adaptations to minimize acoustic interference (Hage et al., 2013; Luo et al., 2015). The dynamics and variability of such strategies are important factors for explaining the high diversity of behavioral adaptations reported in earlier studies. We must keep in mind that the discriminability of a signal from the background is dictated by the difference of the physical parameters between the signal and its background. Essentially, it is unimportant which physical parameter is adjusted, just so the signal has its own physical identity and provides a high level of discriminability from its background. For call-echo assignment, it has been discussed that bats keep an “internal copy” of their broadcasted calls and compare that copy with received echoes (Simmons, 2012). Neural activity occurring before biosonar production in frontal and striatal brain regions could contribute to the formation of such “internal copy” (Weineck et al., 2020). The idea of an “internal copy” is in line with behavioral results showing that correct call-echo assignment is decreased when spectro-temporal properties of the echo are manipulated (Masters and Jacobs, 1989; Masters and Raver, 1996; Masters and Raver, 2000) or when echoes are replaced by noise bursts (Surlykke, 1992). Because of missing behavioral data in *C. perspicillata*, it remains speculative to what extent the echolocation calls need to differ from the playback stimuli so that bats can still extract the signals. When comparing different call parameters against the playback stimuli used in the present study, it becomes clear that some echolocation calls can vastly differ from playback stimuli (S11 Figure). Although having no detection thresholds from *C. perspicillata*, there are some behavioral and electrophysiological results from other bat species that use similar call designs as *C. perspicillata* (for example, *Eptesicus fuscus*: (Chiu et al., 2009; von Stebut and Schmidt, 2001); *Tadarida brasilienisis*: (Bartsch and Schmidt, 1993), *Antrozous pallidus* (Fuzessery, 1994)). Based on these studies, we speculate that *C. perspicillata* can extract signals that differ for one of the following parameters by more than 10 dB in intensity, by at least 0.7 ms in duration, by more than 5 kHz in the peak frequency, by more than 12 kHz in bandwidth, and by more than 6 kHz in the sweep rate from playback stimuli. By considering these thresholds, *C. perspicillata* may be able to extract about 94% of the calls from the playback stimuli. Only 5.96% of the calls did not reach our hypothetical detection thresholds for any of the investigated call parameters. Note that the emission pattern could not be considered for a call-by-call analysis. Thus, it is still probable that the remaining 5.96% of the calls’ echoes could be detected by anticipation of the echo pattern. This ability could be accomplished by grouping the calls (Figure 3A, (Beetz et al., 2018; Beetz et al., 2019)). In the present study, four out of ten bats increased the tendency of grouping the calls, which increases the stimulus rate (Figure 3E).

Neurons of the bat’s auditory midbrain and cortex can likely extract relevant echolocation information when the bats face such high call rates (Bartenstein et al., 2014; Beetz et al., 2018; Beetz et al., 2016a; Beetz et al., 2016b; Beetz et al., 2017; Greiter and Firzlaff, 2017; Macias et al., 2018; Sanderson and Simmons, 2005).

## Acknowledgements

We thank Karen Mesce for her helpful comments on the manuscript.

## References

Adams, A. M., Davis, K. and Smotherman, M. (2017). Suppression of emission rates improves sonar performance by flying bats. Scientific Reports 7.

Amichai, E., Blumrosen, G. and Yovel, Y. (2015). Calling louder and longer: how bats use biosonar under severe acoustic interference from other bats. Proceedings of the Royal Society B-Biological Sciences 282.

Barchi, J. R., Knowles, J. M. and Simmons, J. A. (2013). Spatial memory and stereotypy of flight paths by big brown bats in cluttered surroundings. Journal of Experimental Biology 216, 1053–1063.

Barclay, R. M. R. (1982). Interindividual Use of Echolocation Calls - Eavesdropping by Bats. Behavioral Ecology and Sociobiology 10, 271–275.

Barone, P., Clarey, J. C., Irons, W. A. and Imig, T. J. (1996). Cortical synthesis of azimuth-sensitive single-unit responses with nonmonotonic level tuning: A thalamocortical comparison in the cat. Journal of Neurophysiology 75, 1206–1220.

Bartenstein, S. K., Gerstenberg, N., Vanderelst, D., Peremans, H. and Firzlaff, U. (2014). Echo-acoustic flow dynamically modifies the cortical map of target range in bats. Nature Communications 5, 4668.

Bartsch, E. and Schmidt, S. (1993). Psychophysical Frequency-Modulation Thresholds in a Fm-Bat, Tadarida-Brasiliensis. Hearing Research 67, 128–138.

Bee, M. A. and Micheyl, C. (2008). The cocktail party problem: What is it? How can it be solved? And why should animal behaviorists study it? Journal of Comparative Psychology 122, 235–251.

Beetz, M. J., García-Rosales, F., Kössl, M. and Hechavarría, J. C. (2018). Robustness of cortical and subcortical processing in the presence of natural masking sounds. Sci Rep.

Beetz, M. J., Hechavarria, J. C. and Kossl, M. (2016a). Cortical neurons of bats respond best to echoes from nearest targets when listening to natural biosonar multi-echo streams. Scientific Reports 6.

Beetz, M. J., Hechavarria, J. C. and Kossl, M. (2016b). Temporal tuning in the bat auditory cortex is sharper when studied with natural echolocation sequences. Scientific Reports 6.

Beetz, M. J., Kordes, S., Garcia-Rosales, F., Kossl, M. and Hechavarria, J. C. (2017). Processing of Natural Echolocation Sequences in the Inferior Colliculus of Seba’s Fruit Eating Bat, Carollia perspicillata. Eneuro 4.

Beetz, M. J., Kossl, M. and Hechavarria, J. C. (2019). Adaptations in the call emission pattern of frugivorous bats when orienting under challenging conditions. Journal of Comparative Physiology a-Neuroethology Sensory Neural and Behavioral Physiology 205, 457–467.

Behr, O. and von Helversen, O. (2004). Bat serenades - complex courtship songs of the sac-winged bat (*Saccopteryx bilineata*). Behavioral Ecology and Sociobiology 56, 106–115.

Brumm, H. and Zollinger, S. A. (2011). The evolution of the Lombard effect: 100 years of psychoacoustic research. Behaviour 148, 1173–1198.

Bullock, T. H., Hamstra, R. H. and Scheich, H. (1972). Jamming Avoidance Response of High-Frequency Electric Fish .1. General Features. Journal of Comparative Physiology 77, 1–22.

Chiu, C., Xian, W. and Moss, C. F. (2008). Flying in silence: Echolocating bats cease vocalizing to avoid sonar jamming. Proceedings of the National Academy of Sciences of the United States of America 105, 13116–13121.

Chiu, C., Xian, W. and Moss, C. F. (2009). Adaptive echolocation behavior in bats for the analysis of auditory scenes. Journal of Experimental Biology 212, 1392–1404.

Corcoran, A. J. and Moss, C. F. (2017). Sensing in a noisy world: lessons from auditory specialists, echolocating bats. Journal of Experimental Biology 220, 4554–4566.

Cvikel, N., Levin, E., Hurme, E., Borissov, I., Boonman, A., Amichai, E. and Yovel, Y. (2015). On-board recordings reveal no jamming avoidance in wild bats. Proceedings of the Royal Society B-Biological Sciences 282.

Eckmeier, D., Geurten, B. R. H., Kress, D., Mertes, M., Kern, R., Egelhaaf, M. and Bischof, H. J. (2008). Gaze Strategy in the Free Flying Zebra Finch (Taeniopygia guttata). Plos One 3.

Falk, B., Jakobsen, L., Surlykke, A. and Moss, C. F. (2014). Bats coordinate sonar and flight behavior as they forage in open and cluttered environments. Journal of Experimental Biology 217, 4356–4364.

Fuzessery, Z. M. (1994). Response Selectivity for Multiple Dimensions of Frequency Sweeps in the Pallid Bat Inferior Colliculus. Journal of Neurophysiology 72, 1061–1079.

Ganguly, K. and Kleinfeld, D. (2004). Goal-directed whisking increases phase-locking between vibrissa movement and electrical activity in primary sensory cortex in rat. Proceedings of the National Academy of Sciences of the United States of America 101, 12348–12353.

Gao, L., Balakrishnan, S., He, W. K., Yan, Z. and Müller, R. (2011). Ear Deformations Give Bats a Physical Mechanism for Fast Adaptation of Ultrasonic Beam Patterns. Physical Review Letters 107.

Gillam, E. H. and McCracken, G. F. (2007). Variability in the echolocation of Tadarida brasiliensis: effects of geography and local acoustic environment. Animal Behaviour 74, 277–286.

Gillam, E. H., Ulanovsky, N. and McCracken, G. F. (2007). Rapid jamming avoidance in biosonar. Proc Biol Sci 274, 651–660.

Götze, S., Koblitz, J. C., Denzinger, A. and Schnitzler, H. U. (2016). No evidence for spectral jamming avoidance in echolocation behavior of foraging pipistrelle bats. Scientific Reports 6.

Greiter, W. and Firzlaff, U. (2017). Echo-acoustic flow shapes object representation in spatially complex acoustic scenes. Journal of Neurophysiology, jn 00860 2016.

Griffin, D. R. (1953). Bat Sounds under Natural Conditions, with Evidence for Echolocation of Insect Prey. Journal of Experimental Zoology 123, 435–465.

Habersetzer, J. (1981). Adaptive Echolocation Sounds in the Bat *Rhinopoma-Hardwickei* - a Field-Study. J Comp Physiol 144, 559–566.

Hage, S. R., Jiang, T. L., Berquist, S. W., Feng, J. and Metzner, W. (2013). Ambient noise induces independent shifts in call frequency and amplitude within the Lombard effect in echolocating bats. Proceedings of the National Academy of Sciences of the United States of America 110, 4063–4068.

Hechavarría, J. C. and Kössl, M. (2014). Footprints of inhibition in the response of cortical delay-tuned neurons of bats. Journal of Neurophysiology 111, 1703–1716.

Henson, O. W., Pollak, G. D., Kobler, J. B., Henson, M. M. and Goldman, L. J. (1982). Cochlear Microphonic Potentials Elicited by Biosonar Signals in Flying Bats, *Pteronotus-P-Parnellii*. Hearing Research 7, 127–147.

Hiryu, S., Bates, M. E., Simmons, J. A. and Riquimaroux, H. (2010). FM echolocating bats shift frequencies to avoid broadcast-echo ambiguity in clutter. Proc Natl Acad Sci USA 107, 7048–7053.

Ibanez, C., Juste, J., Lopez-Wilchis, R. and Nunez-Garduno, A. (2004). Habitat variation and jamming avoidance in echolocation calls of the sac-winged bat (*Balantiopteryx plicata*). Journal of Mammalogy 85, 38–42.

Jarvis, J., Jackson, W. and Smotherman, M. (2013). Groups of bats improve sonar efficiency through mutual suppression of pulse emissions. Frontiers in Physiology 4.

Jung, K., Kalko, E. K. V. and von Helversen, O. (2007). Echolocation calls in Central American emballonurid bats: signal design and call frequency alternation. Journal of Zoology 272, 125–137.

Kalko, E. K. V. (1995). Insect Pursuit, Prey Capture and Echolocation in Pipistrelle Bats (Microchiroptera). Animal Behaviour 50, 861–880.

Kalko, E. K. V. and Schnitzler, H. U. (1989). The Echolocation and Hunting Behavior of Daubenton Bat, *Myotis-Daubentoni*. Behavioral Ecology and Sociobiology 24, 225–238.

Knowles, J. M., Barchi, J. R., Gaudette, J. E. and Simmons, J. A. (2015). Effective biosonar echo-to-clutter rejection ratio in a complex dynamic scene. Journal of the Acoustical Society of America 138, 1090–1101.

Kössl, M., Hechavarría, J. C., Voss, C., Macías, S., Mora, E. C. and Vater, M. (2014). Neural maps for target range in the auditory cortex of echolocating bats. Current Opinion in Neurobiology 24, 68–75.

Kothari, N. B., Wohlgemuth, M. J., Hulgard, K., Surlykke, A. and Moss, C. F. (2014). Timing matters: sonar call groups facilitate target localization in bats. Frontiers in Physiology 5.

Land, M. F. (2015). Eye movements of vertebrates and their relation to eye form and function. Journal of Comparative Physiology a-Neuroethology Sensory Neural and Behavioral Physiology 201, 195–214.

Leonard, M. L. and Fenton, M. B. (1984). Echolocation Calls of *Euderma-Maculatum* (Vespertilionidae) - Use in Orientation and Communication. Journal of Mammalogy 65, 122–126.

Levin, E., Roll, U., Dolev, A., Yom-Tov, Y. and Kronfeld-Shcor, N. (2013). Bats of a Gender Flock Together: Sexual Segregation in a Subtropical Bat. Plos One 8.

Lin, Y. and Abaid, N. (2015). Modeling perspectives on echolocation strategies inspired by bats flying in groups. Journal of Theoretical Biology 387, 46–53.

Luo, J. H., Goerlitz, H. R., Brumm, H. and Wiegrebe, L. (2015). Linking the sender to the receiver: vocal adjustments by bats to maintain signal detection in noise. Scientific Reports 5.

Macias, S., Luo, J. H. and Moss, C. F. (2018). Natural echolocation sequences evoke echo-delay selectivity in the auditory midbrain of the FM bat, Eptesicus fuscus. Journal of Neurophysiology 120, 1323–1339.

Macias, S., Mora, E. C., Hechavarria, J. C. and Kossl, M. (2016). Echo-level compensation and delay tuning in the auditory cortex of the mustached bat. European Journal of Neuroscience 43, 1647–1660.

Masters, W. M. and Jacobs, S. C. (1989). Target Detection and Range Resolution by the Big Brown Bat (Eptesicus-Fuscus) Using Normal and Time-Reversed Model Echoes. Journal of Comparative Physiology a-Sensory Neural and Behavioral Physiology 166, 65–73.

Masters, W. M. and Raver, K. A. S. (1996). The degradation of distance discrimination in big brown bats (Eptesicus fuscus) caused by different interference signals. Journal of Comparative Physiology a-Sensory Neural and Behavioral Physiology 179, 703–713.

Masters, W. M. and Raver, K. A. S. (2000). Range discrimination by big brown bats (Eptesicus fuscus) using altered model echoes: Implications for signal processing. Journal of the Acoustical Society of America 107, 625–637.

Mazar, O. and Yovel, Y. (2019). A spectral jamming avoidance response does not help bats deal with jamming. bioRxiv, 2019.12.16.876086.

Miller, L. A. and Degn, H. J. (1981). The Acoustic Behavior of 4 Species of Vespertilionid Bats Studied in the Field. Journal of Comparative Physiology 142, 67–74.

Moss, C. F. and Surlykke, A. (2010). Probing the natural scene by echolocation in bats. Frontiers in Behavioral Neuroscience 4.

Obrist, M. K. (1995). Flexible Bat Echolocation - the Influence of Individual, Habitat and Conspecifics on Sonar Signal-Design. Behavioral Ecology and Sociobiology 36, 207–219.

Park, T. J. and Pollak, G. D. (1993). Gaba Shapes a Topographic Organization of Response Latency in the Moustache Bats Inferior Colliculus. Journal of Neuroscience 13, 5172–5187.

Parsons, K. N., Jones, G. and Greenaway, F. (2003). Swarming activity of temperate zone microchiropteran bats: effects of season, time of night and weather conditions. Journal of Zoology 261, 257–264.

Petrites, A. E., Eng, O. S., Mowlds, D. S., Simmons, J. A. and DeLong, C. M. (2009). Interpulse interval modulation by echolocating big brown bats (*Eptesicus fuscus*) in different densities of obstacle clutter. J Comp Physiol A 195, 603–617.

Ratcliffe, J. M., ter Hofstede, H. M., Avila-Flores, R., Fenton, M. B., McCracken, G. F., Biscardi, S., Blasko, J., Gillam, E., Orprecio, J. and Spanjer, G. (2004). Conspecifics influence call design in the Brazilian free-tailed bat, Tadarida brasiliensis. Canadian Journal of Zoology-Revue Canadienne De Zoologie 82, 966–971.

Ribak, G., Egge, A. R. and Swallow, J. G. (2009). Saccadic head rotations during walking in the stalk-eyed fly (Cyrtodiopsis dalmanni). Proceedings of the Royal Society B-Biological Sciences 276, 1643–1649.

Roverud, R. C. and Grinnell, A. D. (1985a). Discrimination Performance and Echolocation Signal Integration Requirements for Target Detection and Distance Determination in the CF FM Bat, *Noctilio-Albiventris*. J Comp Physiol A 156, 447–456.

Roverud, R. C. and Grinnell, A. D. (1985b). Echolocation Sound Features Processed to Provide Distance Information in the Cf Fm Bat, Noctilio-Albiventris - Evidence for a Gated Time Window Utilizing Both Cf and Fm Components. Journal of Comparative Physiology a-Sensory Neural and Behavioral Physiology 156, 457–469.

Sanderson, M. I. and Simmons, J. A. (2005). Target representation of naturalistic echolocation sequences in single unit responses from the inferior colliculus of big brown bats. Journal of the Acoustical Society of America 118, 3352–3361.

Schnitzler, H. U., Kalko, E., Miller, L. and Surlykke, A. (1987). The Echolocation and Hunting Behavior of the Bat, Pipistrellus-Kuhli. Journal of Comparative Physiology a-Sensory Neural and Behavioral Physiology 161, 267–274.

Schroeder, C. E., Wilson, D. A., Radman, T., Scharfman, H. and Lakatos, P. (2010). Dynamics of Active Sensing and perceptual selection. Current Opinion in Neurobiology 20, 172–176.

Simmons, J. A. (2012). Bats use a neuronally implemented computational acoustic model to form sonar images. Current Opinion in Neurobiology 22, 311–319.

Simmons, J. A. (2017). Noise interference with echo delay discrimination in bat biosonar. Journal of the Acoustical Society of America 142, 2942–2952.

Simmons, J. A., Fenton, M. B. and Ofarrell, M. J. (1979). Echolocation and Pursuit of Prey by Bats. Science 203, 16–21.

Simmons, J. A., Howell, D. J. and Suga, N. (1975). Information-Content of Bat Sonar Echoes. American Scientist 63, 204–215.

Simmons, J. A., Lavender, W. A., Lavender, B. A., Childs, J. E., Hulebak, K., Rigden, M. R., Sherman, J., Woolman, B. and Ofarrell, M. J. (1978). Echolocation by Free-Tailed Bats (Tadarida). Journal of Comparative Physiology 125, 291–299.

Suga, N. and Manabe, T. (1982). Neural Basis of Amplitude-Spectrum Representation in Auditory-Cortex of the Mustached Bat. Journal of Neurophysiology 47, 225–255.

Suga, N., O’Neill, W. E., Kujirai, K. and Manabe, T. (1983). Specificity of Combination-Sensitive Neurons for Processing of Complex Biosonar Signals in Auditory-Cortex of the Mustached Bat. Journal of Neurophysiology 49, 1573–1626.

Surlykke, A. (1992). Target Ranging and the Role of Time-Frequency Structure of Synthetic Echoes in Big Brown Bats, Eptesicus-Fuscus. Journal of Comparative Physiology a-Sensory Neural and Behavioral Physiology 170, 83–92.

Surlykke, A. and Moss, C. F. (2000). Echolocation behavior of big brown bats, Eptesicus fuscus, in the field and the laboratory. Journal of the Acoustical Society of America 108, 2419–2429.

Takahashi, E., Hyomoto, K., Riquimaroux, H., Watanabe, Y., Ohta, T. and Hiryu, S. (2014). Adaptive changes in echolocation sounds by *Pipistrellus abramus* in response to artificial jamming sounds. Journal of Experimental Biology 217, 2885–2891.

Tarsitano, M. S. and Andrew, R. (1999). Scanning and route selection in the jumping spider Portia labiata. Animal Behaviour 58, 255–265.

Thies, W., Kalko, E. K. V. and Schnitzler, H. U. (1998). The roles of echolocation and olfaction in two Neotropical fruit-eating bats, *Carollia perspicillata* and *C-castanea*, feeding on Piper. Behavioral Ecology and Sociobiology 42, 397–409.

Towal, R. B. and Hartmann, M. J. (2006). Right-left asymmetries in the whisking behavior of rats anticipate head movements. Journal of Neuroscience 26, 8838–8846.

Tressler, J. and Smotherman, M. S. (2009). Context-dependent effects of noise on echolocation pulse characteristics in free-tailed bats. J Comp Physiol A 195, 923–934.

Ulanovsky, N., Fenton, M. B., Tsoar, A. and Korine, C. (2004). Dynamics of jamming avoidance in echolocating bats. Proceedings of the Royal Society B-Biological Sciences 271, 1467–1475.

Ulanovsky, N. and Moss, C. F. (2008). What the bat’s voice tells the bat’s brain. Proc Natl Acad Sci USA 105, 8491–8498.

von Stebut, B. and Schmidt, S. (2001). Frequency discrimination threshold at search call frequencies in the echolocating bat, Eptesicus fuscus. Journal of Comparative Physiology a-Sensory Neural and Behavioral Physiology 187, 287–291.

Watanabe, A. and Takeda, K. (1963). The Change of Discharge Frequency by A.C. Stimulus in a Weak Electric Fish. Journal of Experimental Biology 40, 57-+.

Weineck, K., Garcia-Rosales, F. and Hechavarria, J. C. (2020). Neural oscillations in the fronto-striatal network predict vocal output in bats. Plos Biology 18.

Wheeler, A. R., Fulton, K. A., Gaudette, J. E., Simmons, R. A., Matsuo, I. and Simmons, J. A. (2016). Echolocating Big Brown Bats, *Eptesicus fuscus*, Modulate Pulse Intervals to Overcome Range Ambiguity in Cluttered Surroundings. Frontiers in Behavioral Neuroscience 10.

Wohlgemuth, M. J., Kothari, N. B. and Moss, C. F. (2016a). Action Enhances Acoustic Cues for 3-D Target Localization by Echolocating Bats. Plos Biology 14.

Wohlgemuth, M. J., Luo, J. and Moss, C. F. (2016b). Three-dimensional auditory localization in the echolocating bat. Current Opinion in Neurobiology 41, 78–86.

Yang, L. C., Pollak, G. D. and Resler, C. (1992). Gabaergic Circuits Sharpen Tuning Curves and Modify Response Properties in the Moustache Bat Inferior Colliculus. Journal of Neurophysiology 68, 1760–1774.

Yovel, Y., Melcon, M. L., Franz, M. O., Denzinger, A. and Schnitzler, H. U. (2009). The Voice of Bats: How Greater Mouse-eared Bats Recognize Individuals Based on Their Echolocation Calls. Plos Computational Biology 5.

