## Supplement for "Bats dynamically change echolocation parameters in response to acoustic playback"

**Running Title: Adaptations in response to acoustic playback**

M. Jerome Beetz (*)[1,2], Manfred Kössl[1], Julio C. Hechavarría[1]

**Affiliations**

[1]Institut für Zellbiologie und Neurowissenschaft, Goethe-Universität, Frankfurt/M., Germany

[2] current address: Zoology II Emmy-Noether Animal Navigation Group, Biocenter, University of Würzburg, Germany

* Corresponding author

***Mailing address*:**

M. Jerome Beetz

Zoology II, University of Wuerzburg, Am Hubland, 97074, Wuerzburg, Germany, Tel.: +49 9313184528

**Keywords** *echolocation, active sensing, bioacoustics, signal interference, spatial orientation, Jamming Avoidance Response*

### Supporting information

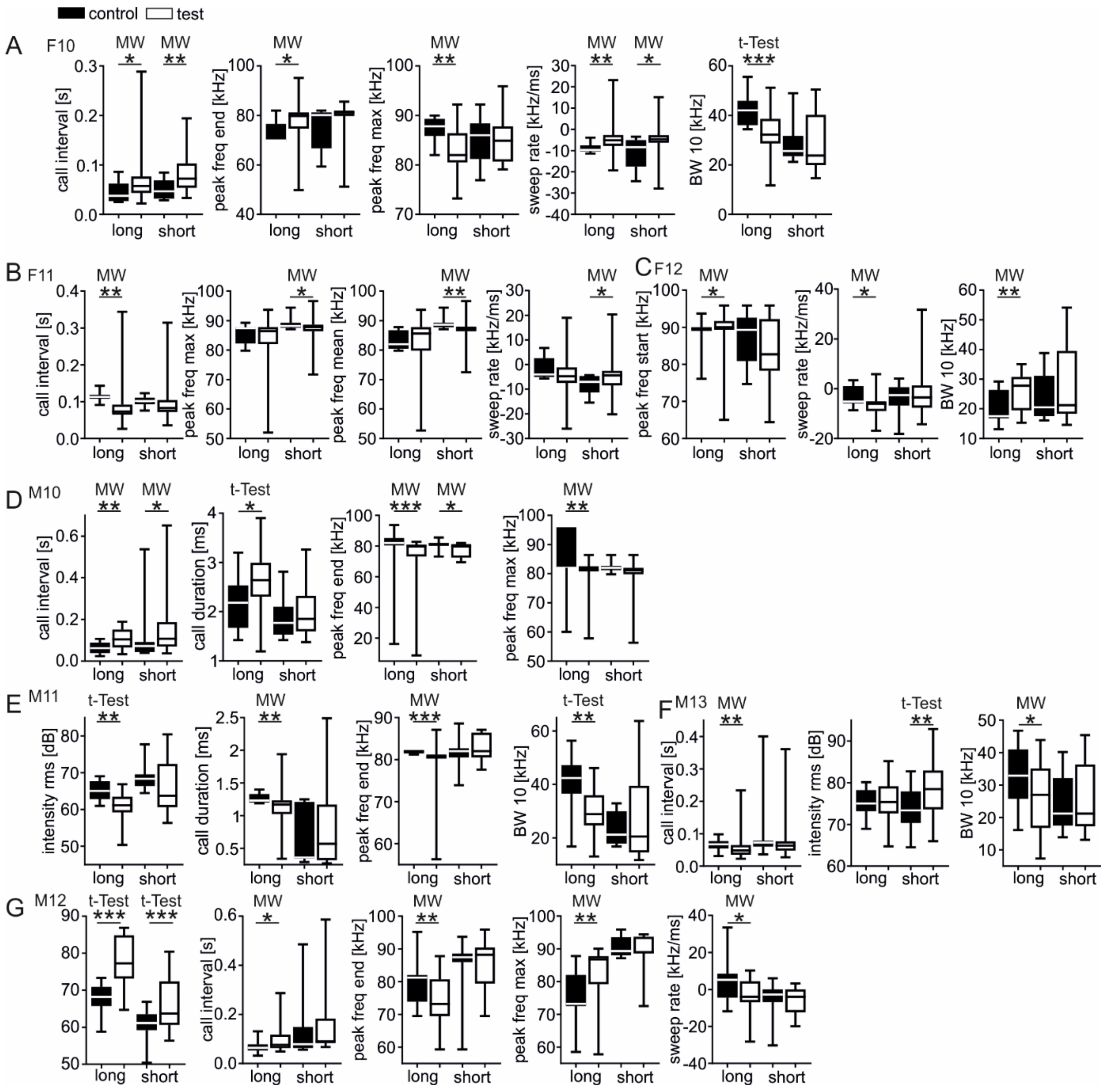


**Figure S1 Individual specific call adaptations in response to playback stimuli**

(A-G) Boxplots from seven individuals showing the call parameters that the bats changed in response to the playback stimulus. Calls recorded under control conditions (absence of playback stimulus) are indicated by black boxplots and white boxplots represent calls recorded under test conditions (presence of playback stimulus). Note that each bat changed different call parameters under test conditions. *BW* = bandwidth; *freq* = frequency; *MW* = mann-whitney test; * $p < 0.05$; ** $p < 0.01$; *** $p < 0.0001$.

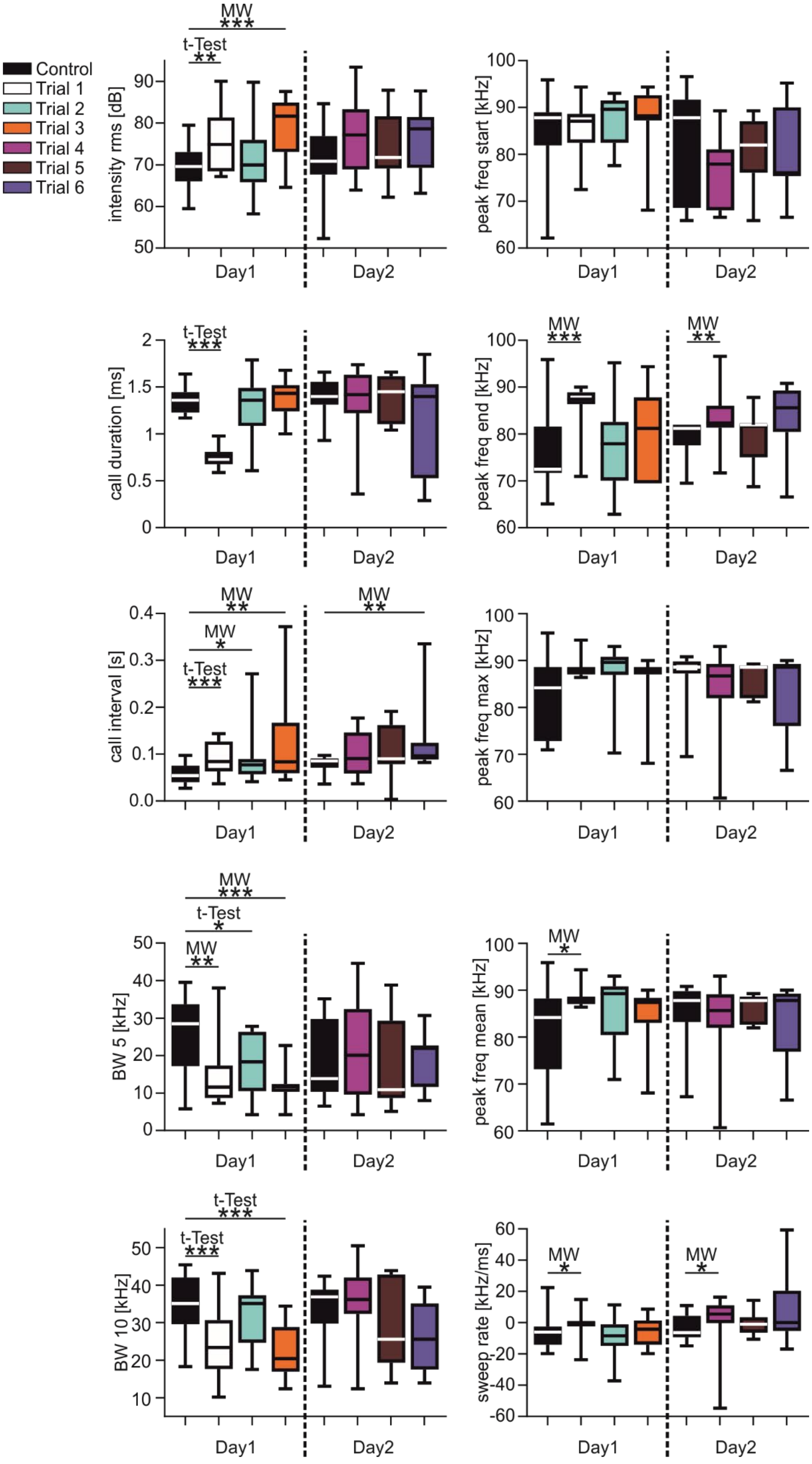


**Figure S2 Bats switch adaptation strategies across trials and days**

Call parameters are shown as boxplots for each trial (6 test trials and 2 control trials) across two days from one bat (F8). For visualization, each trial is color coded. *BW* = bandwidth; *freq* = frequency; * p < 0.05; ** p < 0.01; *** p < 0.0001

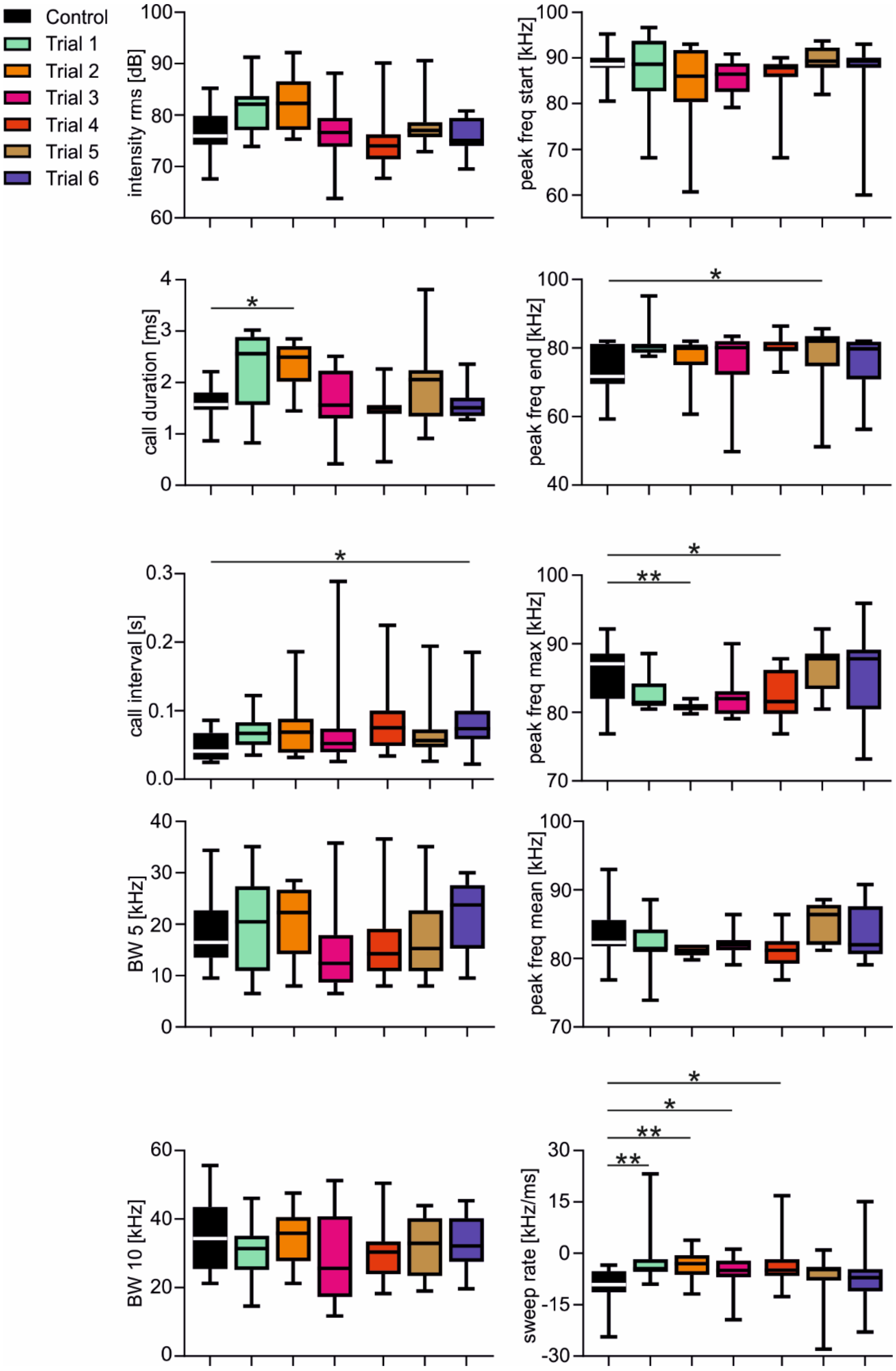

**Figure S3 Bats switch adaptation strategies across trials**

Call parameters are shown as boxplots for each trial (6 test trials and 1 control trial) from one bat (F10). For visualization, each trial is color coded. *BW* = bandwidth; *freq* = frequency; $p < 0.05$; ** $p < 0.01$

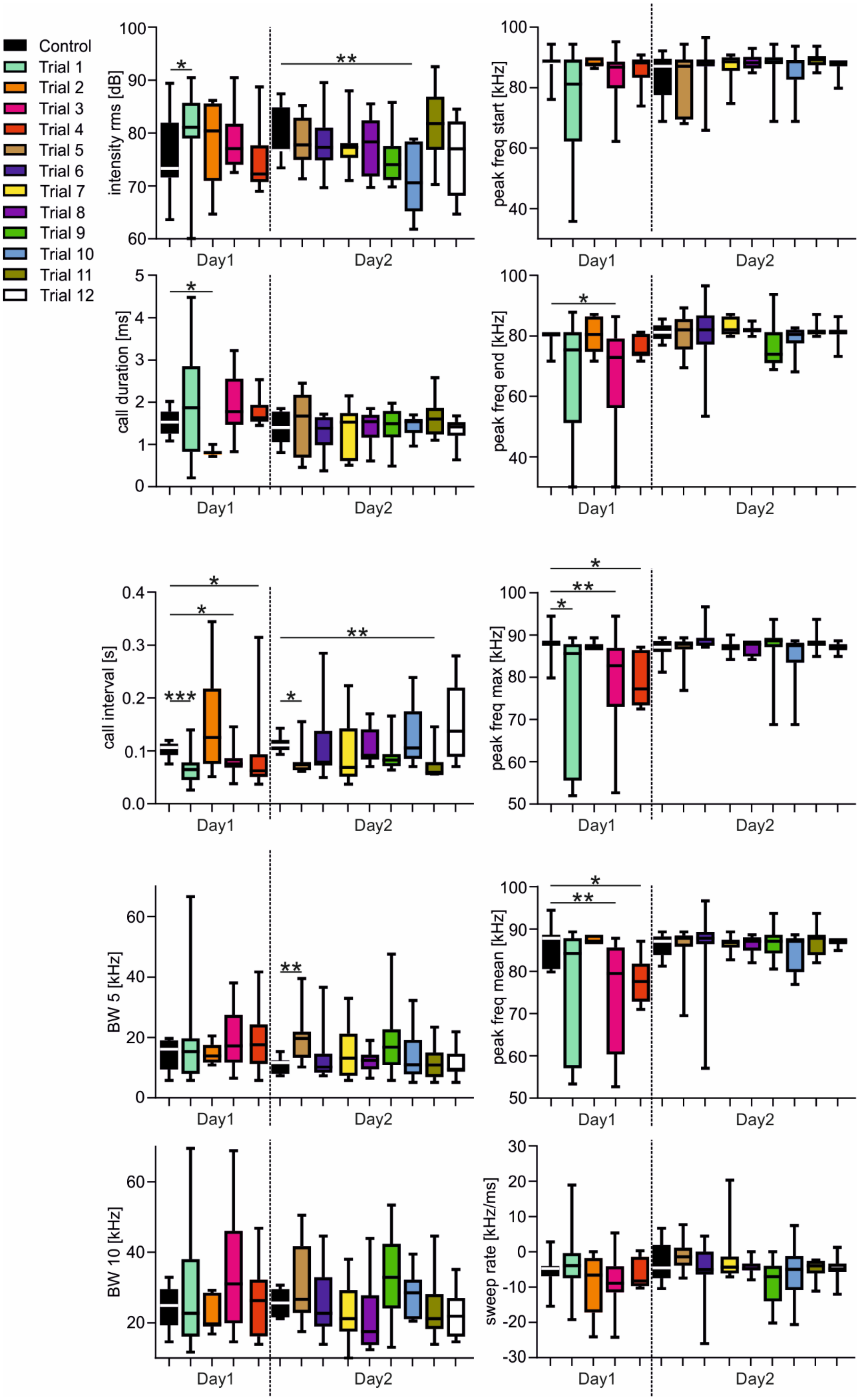


**Figure S4 Bats switch adaptation strategies across trials and days**

Call parameters are shown as boxplots for each trial (12 test trials and 2 control trials) across two days from one bat (F11). For visualization, each trial is color coded. *BW* = bandwidth; *freq* = frequency; * p < 0.05; ** p < 0.01; *** p < 0.0005

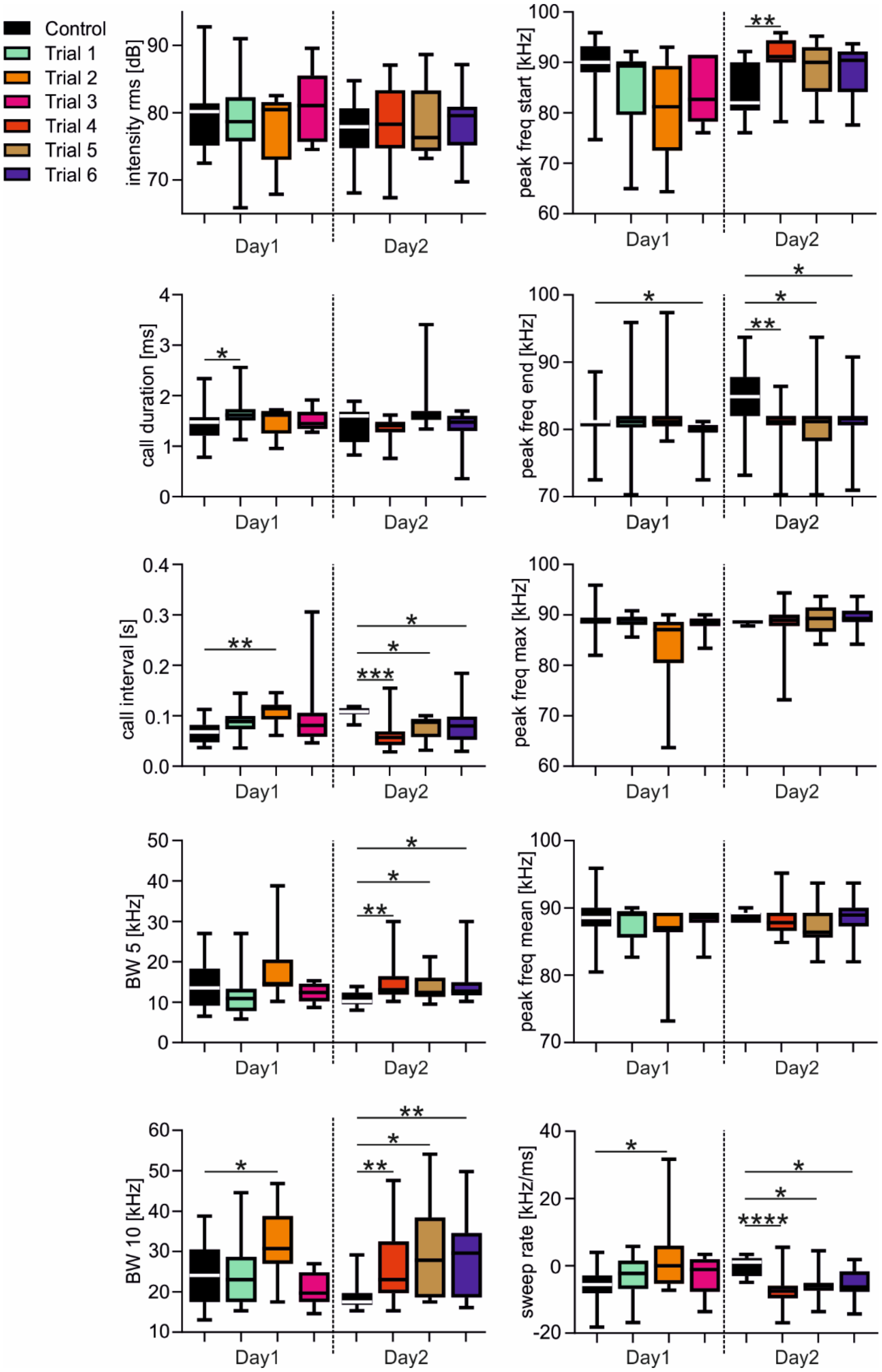


**Figure S5 Bats switch adaptation strategies across trials and days**

Call parameters are shown as boxplots for each trial (6 test trials and 2 control trials) across two days from one bat (F12). For visualization, each trial is color coded. *BW* = bandwidth; *freq* = frequency; * $p < 0.05$; ** $p < 0.01$; *** $p < 0.0005$, **** $p < 0.0001$

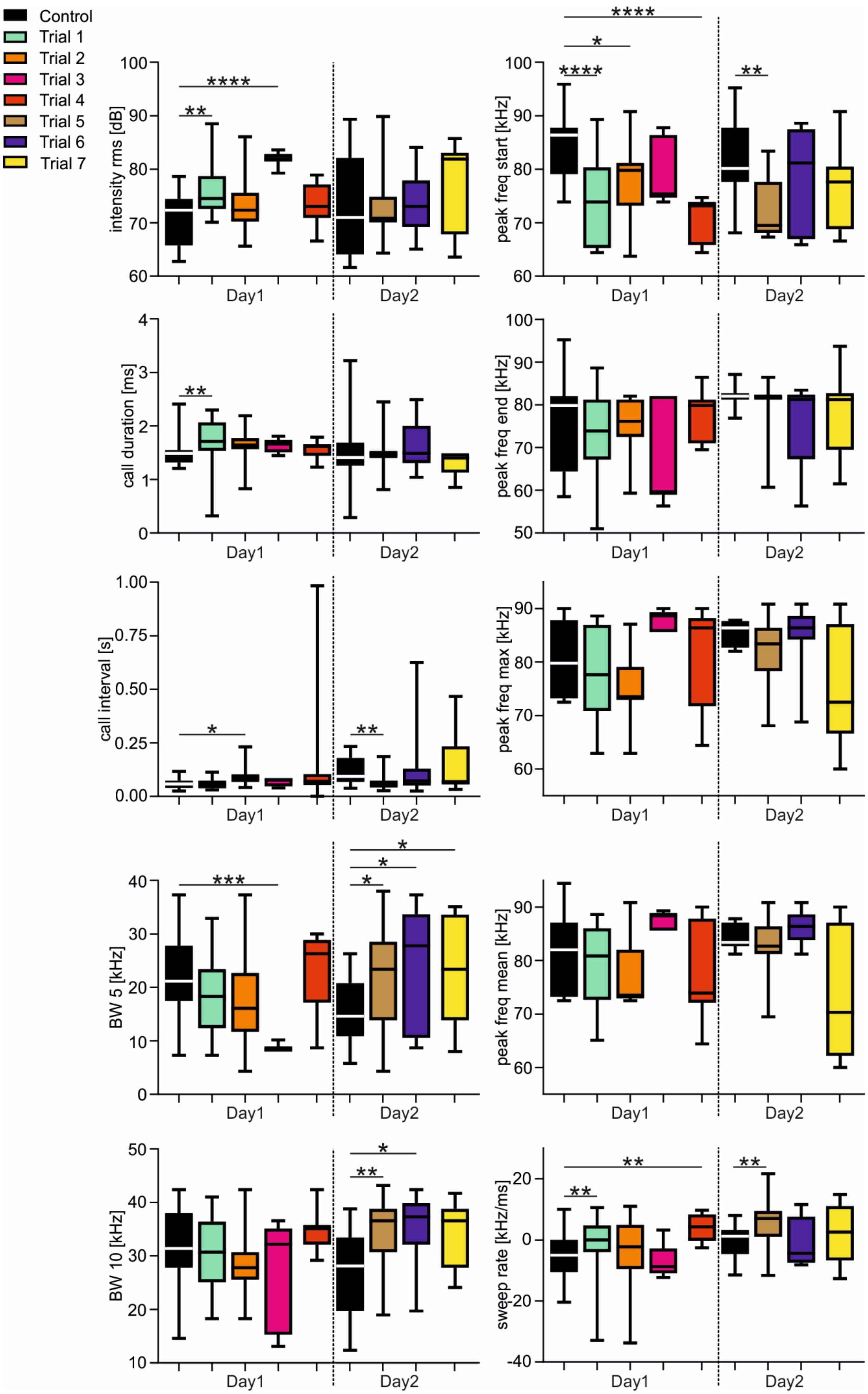


**Figure S6 Bats switch adaptation strategies across trials and days**

Call parameters are shown as boxplots for each trial (7 test trials and 2 control trials) across two days from one bat (M9). For visualization, each trial is color coded. *BW* = bandwidth; *freq* = frequency; * $p < 0.05$; ** $p < 0.01$; *** $p < 0.0005$, **** $p < 0.0001$

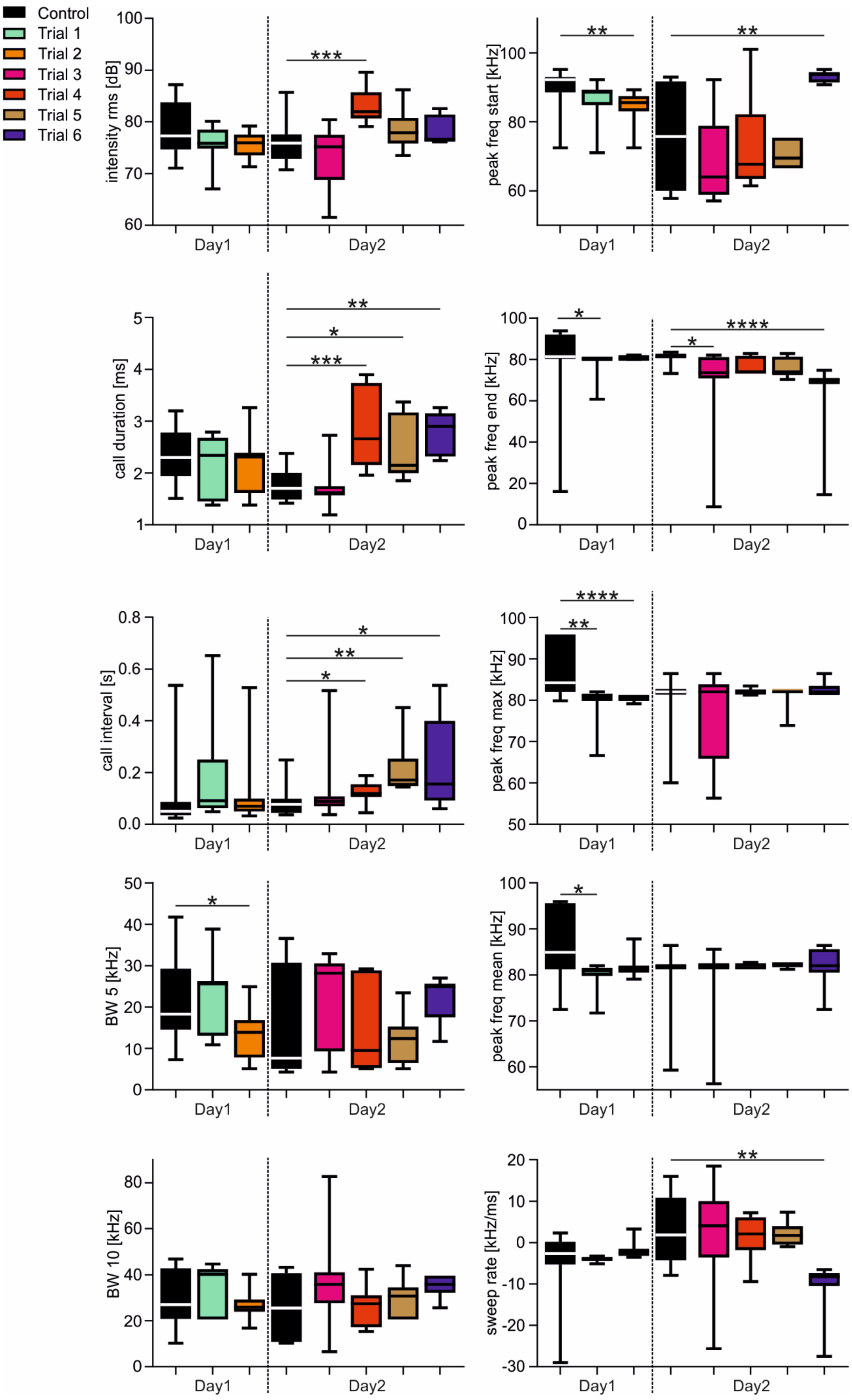

Control
Trial 1
Trial 2
Trial 3
Trial 4
Trial 5
Trial 6
intensity rms [dB]
peak freq start [kHz]
call duration [ms]
peak freq end [kHz]
call interval [s]
peak freq max [kHz]
BW 5 [kHz]
peak freq mean [kHz]
BW 10 [kHz]
sweep rate [kHz/ms]
Day1
Day2

**Figure S7 Bats switch adaptation strategies across trials and days**

Call parameters are shown as boxplots for each trial (6 test trials and 2 control trials) across two days from one bat (M10). For visualization, each trial is color coded. *BW* = bandwidth; *freq* = frequency; * p < 0.05; ** p < 0.01; *** p < 0.0005, **** p < 0.0001

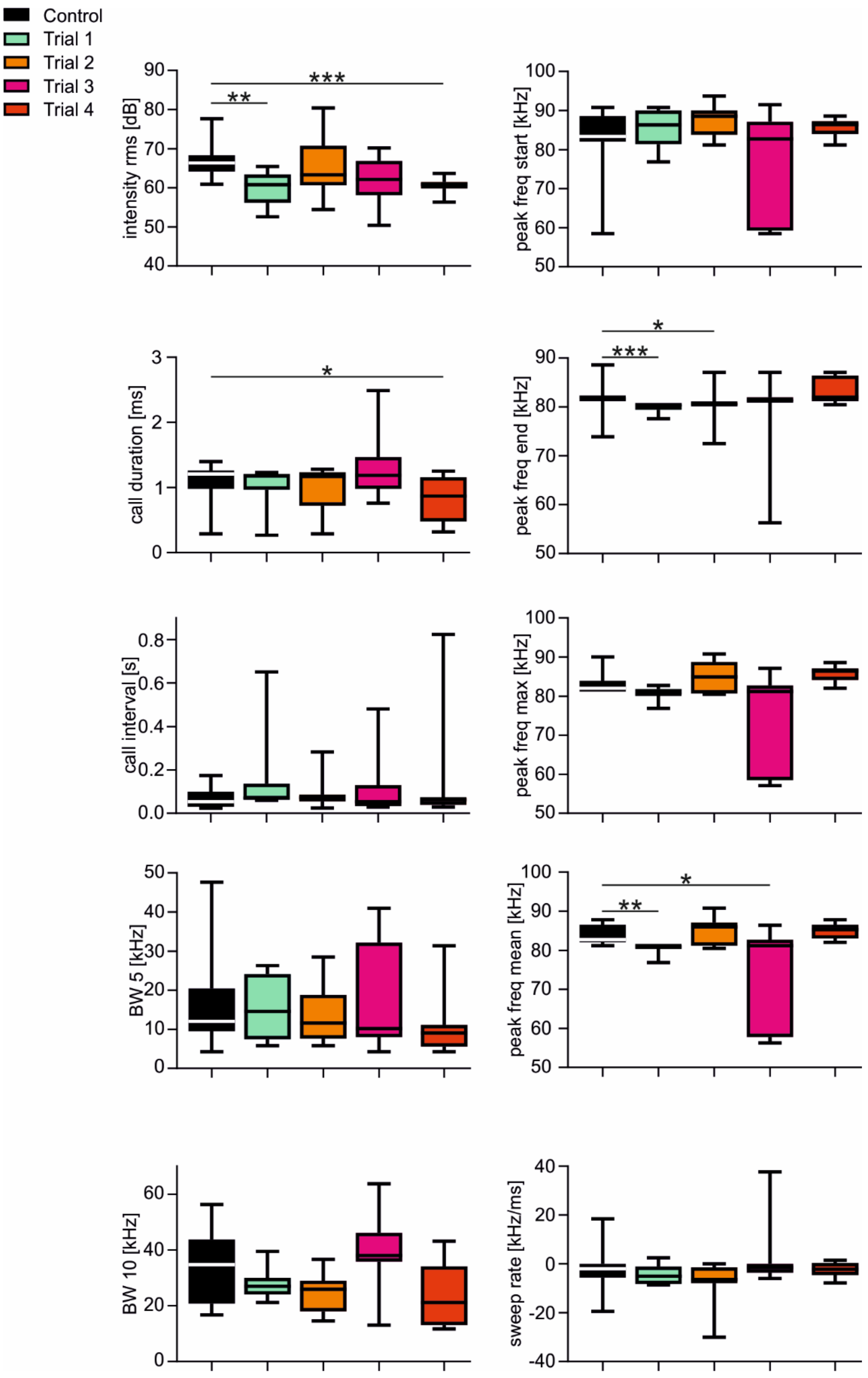

**Figure S8 Bats switch adaptation strategies across trials and days**

Call parameters are shown as boxplots for each trial (4 test trials and 2 control trials) from one bat (M11). For visualization, each trial is color coded. *BW* = bandwidth; *freq* = frequency; * p < 0.05; ** p < 0.01; *** p < 0.0005

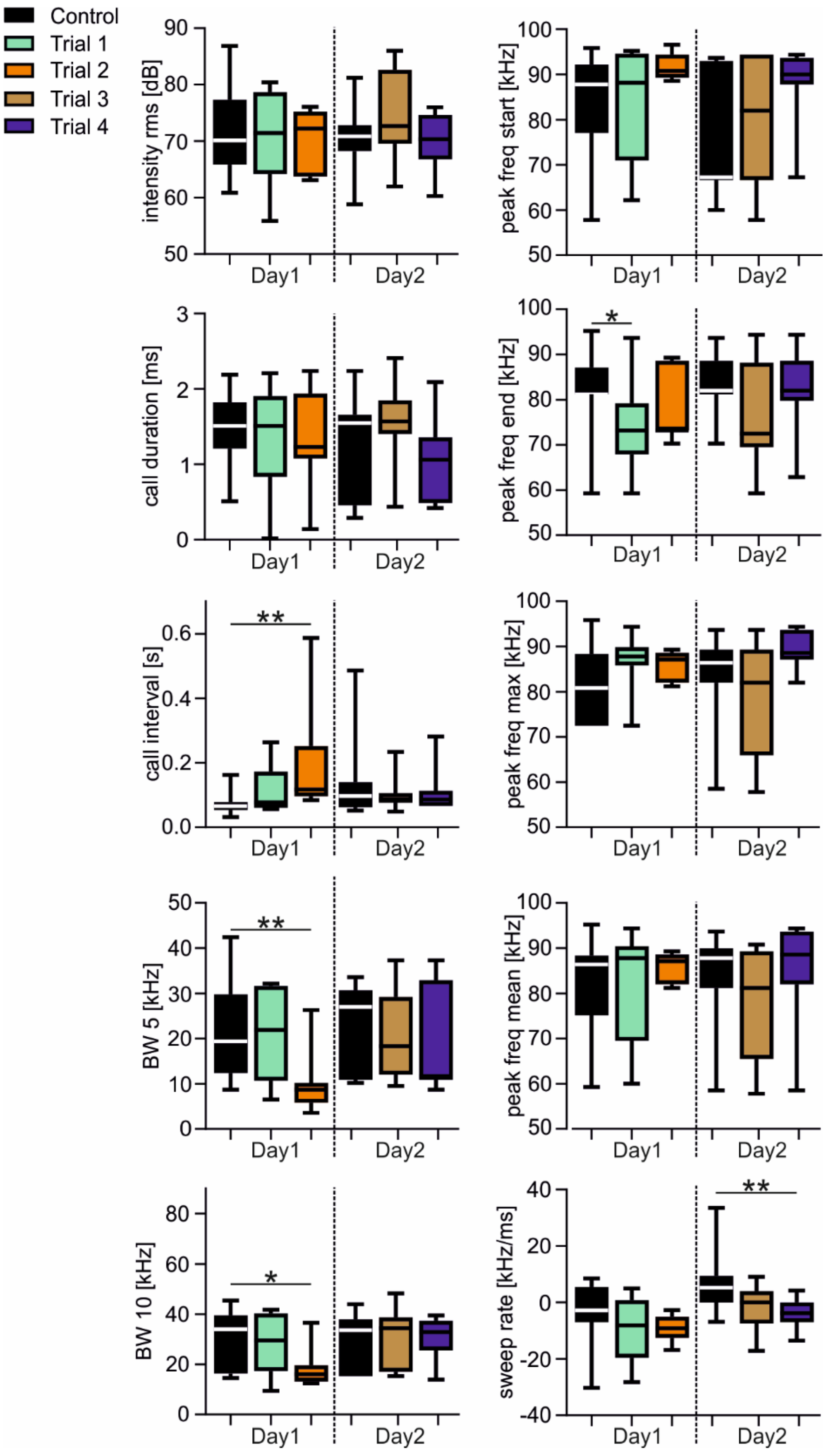


**Figure S9 Bats switch adaptation strategies across trials and days**

Call parameters are shown as boxplots for each trial (6 test trials and 2 control trials) across two days from one bat (M12). For visualization, each trial is color coded. *BW* = bandwidth; *freq* = frequency; * p < 0.05; ** p < 0.01

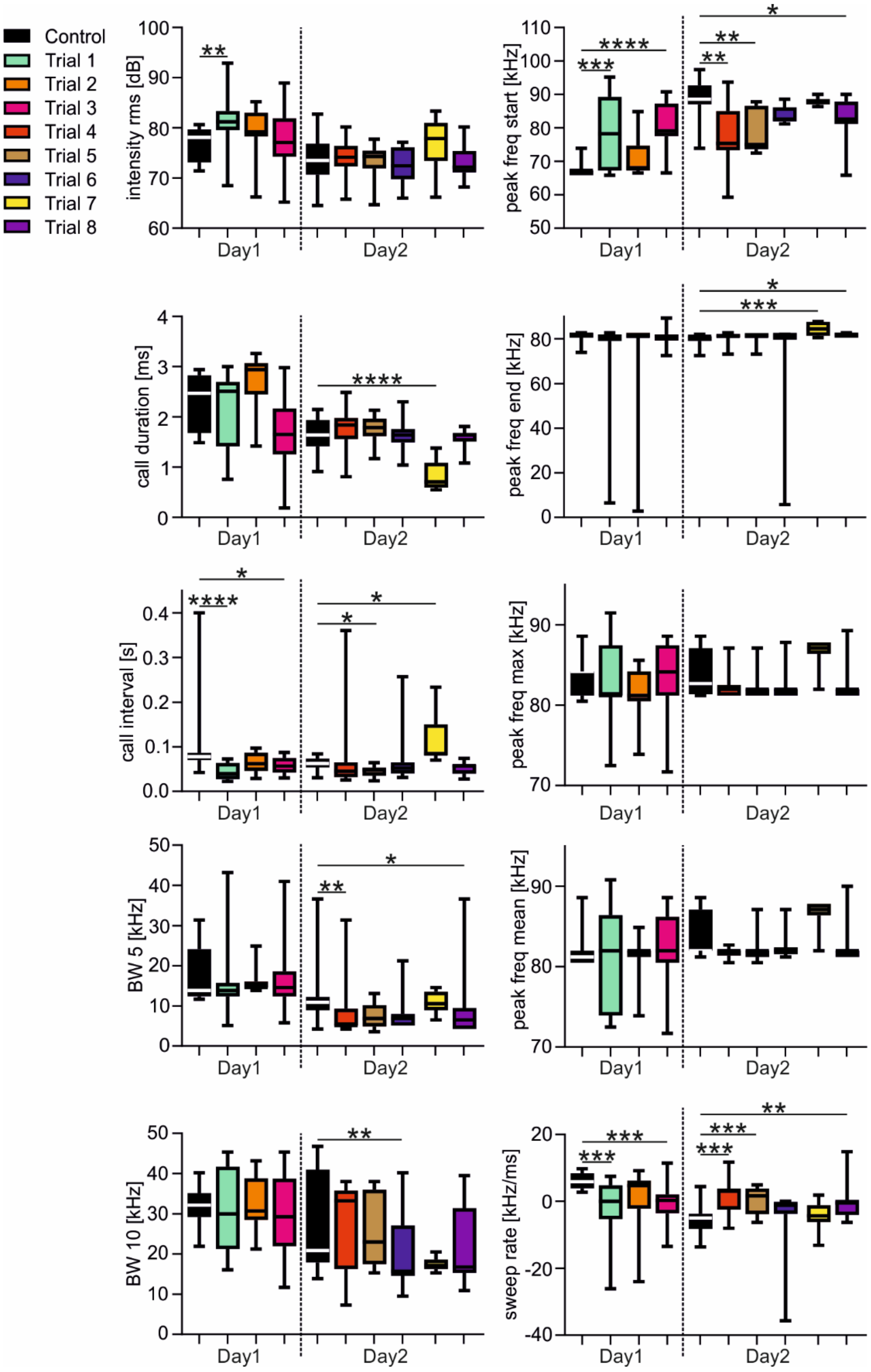


**Figure S10 Bats switch adaptation strategies across trials and days**

Call parameters are shown as boxplots for each trial (8 test trials and 2 control trials) across two days from one bat (M13). For visualization, each trial is color coded. *BW* = bandwidth; *freq* = frequency; * $p < 0.05$; ** $p < 0.01$; *** $p < 0.0005$, **** $p < 0.0001$

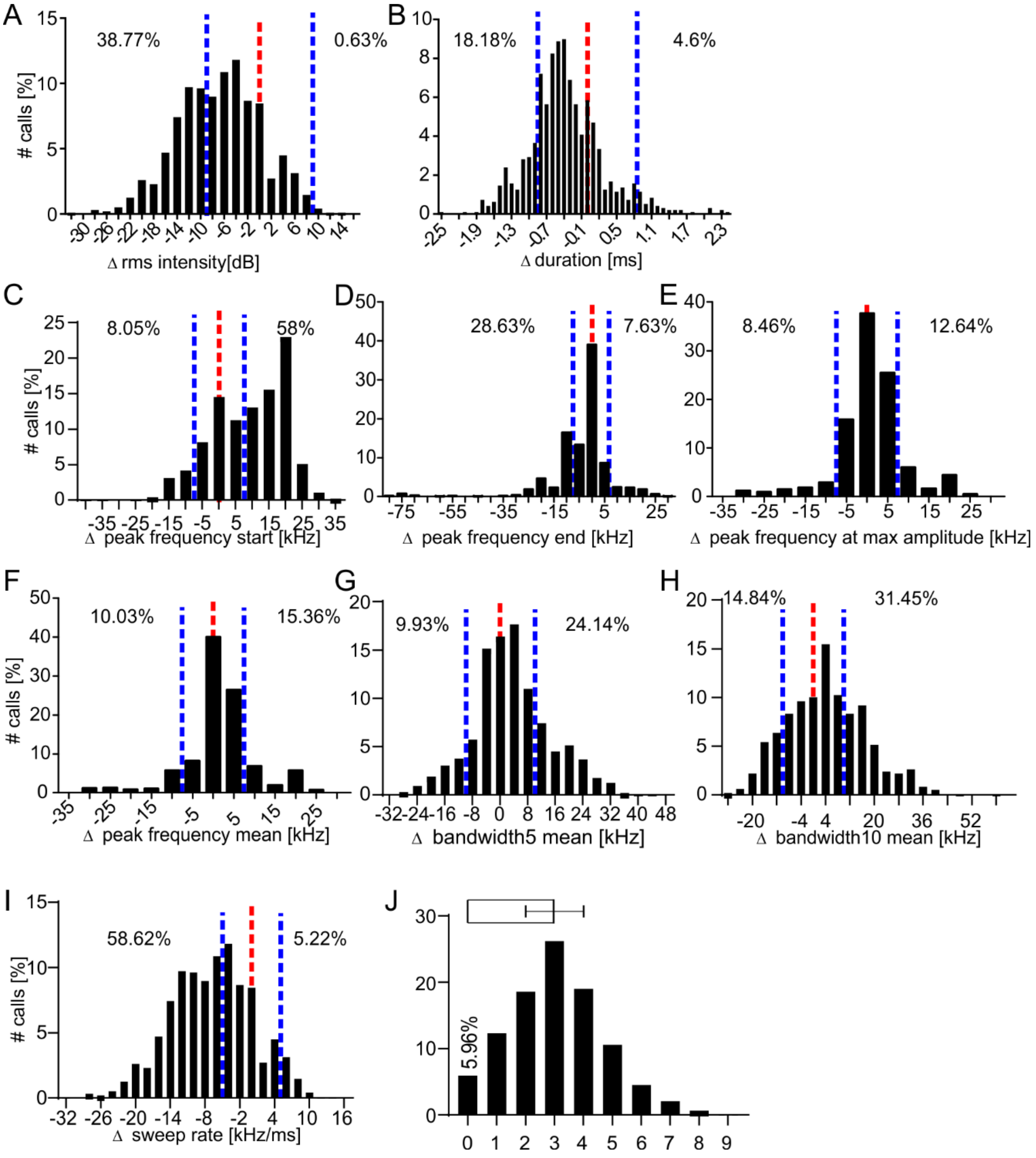


**Figure S11 Comparison between emitted echolocation calls and play backed calls**

**(A-I)** Histrograms demonstrating the differences of the echolocation calls from the playback calls for different call parameters (intensity **(A)**, duration **(B)**, peak frequency start **(C)**, peak frequency end **(D)**, peak frequency max **(E)**, peak frequency mean **(F)**, bandwidth 5 **(G)**, bandwidth 10 **(H)**, sweep rate **(I)**. Red vertical dashed lines indicate the zero line. Calls at the zero line were equal to the playback stimulus with respect to the call parameter. Blue vertical dashed lines indicate lower and upper thresholds for defining the calls to be different from the playback calls. For each tested call parameter, the percentage of calls that are below or above the thresholds are indicated in each plot.

**(J)** Histogram summarizing how many call parameters were different from the control call of the bat that was recorded beforehand and used as interfering playback. Only 5.96% of the calls had a call design similar to the control call. A summary bar with standard deviation is shown on top of the histogram.

Supple Table 1 P-values from the statistics comparing call parameters across subsequent pendulum swings (control trials) from nine bats. Across subsequent control trials, bats did not change call duration, intensity, starting peak frequency (P F start), sweep rate, terminal peak frequency (P F end), maximum peak frequency (P F max), mean peak frequency (P F mean), bandwidth 5 (BW 5), and bandwidth 10 (BW10) (indicated by blue cells). Four bats (F9, F10, F11, and M12) varied significantly the call intervals across subsequent trials (indicated by red cells). Namely, they increased the call intervals which may indicate a habituation to the pendulum. For **F8, F11, F12, M9, M10, M11,** and **M12**, p-values result either from an one-way ANOVA (if data follow a Gaussian distribution based on D´Agostino-Pearson omnibus normality test) or from a Kruskal-Wallis test (if data do not follow a Gaussian distribution). For **F9** and **F10**, p-values result either from an unpaired t-test (if data follow a Gaussian distribution based on D´Agostino-Pearson omnibus normality test) or from a Mann-Whitney test (if data do not follow a Gaussian distribution).

| **Animal ID** | **F8** | **F9** | **F10** | **F11** | **F12** | **M9** | **M10** | **M11** | **M12** |
|---|---|---|---|---|---|---|---|---|---|
| **N calls/ trial** | 19,17,14 | 20,12 | 25,11 | 20,14,17,19 | 12,17,14 | 24,21,14 | 15,15,18 | 18,16,26 | 20,13,12 |
| **Interval** | 0.63 | 0.002 | 0.0002 | 0.004 | 0.5 | 0.09 | 0.9 | 0.5 | 0.0012 |
| **Duration** | 0.17 | 0.26 | 0.37 | 0.15 | 0.24 | 0.66 | 0.64 | 0.11 | 0.06 |
| **Intensity** | 0.53 | 0.83 | 0.17 | 0.31 | 0.26 | 0.08 | 0.1 | 0.16 | 0.06 |
| **P F start** | 0.1 | 0.13 | 0.57 | 0.12 | 0.21 | 0.6 | 0.11 | 0.12 | 0.81 |
| **Sweep rate** | 0.27 | 0.96 | 0.89 | 0.39 | 0.39 | 0.97 | 0.07 | 0.23 | 0.21 |
| **P F end** | 0.96 | 0.18 | 0.59 | 0.8 | 0.31 | 0.66 | 0.49 | 0.39 | 0.72 |
| **P F max** | 0.13 | 0.32 | 0.95 | 0.1 | 0.57 | 0.46 | 0.05 | 0.06 | 0.71 |
| **P F mean** | 0.08 | 0.73 | 0.31 | 0.18 | 0.3 | 0.78 | 0.09 | 0.38 | 0.42 |
| **BW5** | 0.07 | 0.81 | 0.07 | 0.1 | 0.07 | 0.94 | 0.47 | 0.12 | 0.37 |
| **BW10** | 0.32 | 0.78 | 0.15 | 0.46 | 0.4 | 0.69 | 0.84 | 0.62 | 0.58 |

Supple Table 2 Changes of the call parameters induced by the presence of playback stimuli. Calls were grouped into long (*l*) and short (*s*) delay calls. In contrast to table 2, supple table 2 indicate the exact p-values in its cells. Blue and red cells indicated non-significant ($p > 0.05$) and significant ($p < 0.05$) differences. p-values result either from an one-way ANOVA (if data follow a Gaussian distribution based on D´Agostino-Pearson omnibus normality test) or from a Kruskal-Wallis test (if data do not follow a Gaussian distribution). F = female; M = male; l = long delay calls; p f = peak frequency; s = short delay calls

| **Animal ID (n calls control/n calls test)** | **F8 (21/34;20/48)** | | **F9 (33/95;13/51)** | | **F10 (10/64; 12/28)** | | **F11 (12/96; 10/62)** | | **F12 (12/46; 17/47)** | | **M9 (30/57; 17/37)** | | **M10 (22/32; 13/27)** | | **M11 (13/30; 9/11)** | | **M12 (19/30; 12/22)** | | **M13** | |
|---|---|---|---|---|---|---|---|---|---|---|---|---|---|---|---|---|---|---|---|---|
| **Delay** | **l** | **s** | **l** | **s** | **l** | **s** | **l** | **s** | **l** | **s** | **l** | **s** | **l** | **s** | **l** | **s** | **l** | **s** | **l** | **s** |
| **Interval** | 0.002 | 0.0002 | 0.24 | 0.77 | 0.03 | 0.008 | 0.002 | 0.07 | 0.19 | 0.76 | 0.56 | 0.31 | 0.003 | 0.05 | 0.22 | 0.2 | 0.03 | 0.17 | 0.006 | 0.18 |
| **Duration** | 0.02 | 0.68 | $<1x10^{-4}$ | 0.001 | 0.54 | 0.45 | 0.73 | 0.53 | 0.59 | 0.28 | 0.03 | 0.4 | 0.01 | 0.51 | 0.004 | 0.97 | 0.83 | 0.62 | 0.35 | 0.16 |
| **Intensity** | 0.06 | 0.0006 | 0.17 | 0.003 | 0.06 | 0.55 | 0.07 | 0.3 | 0.98 | 0.47 | 0.002 | 0.84 | 0.11 | 0.33 | 0.001 | 0.32 | $<1x10^{-4}$ | 0.0007 | 0.97 | 0.03 |
| **P F start** | 0.93 | 0.06 | 0.03 | $<1x10^{-4}$ | 0.53 | 0.25 | 0.51 | 0.33 | 0.04 | 0.62 | $<1x10^{-4}$ | 0.06 | 0.31 | 0.07 | 0.43 | >0.99 | 0.74 | 0.21 | 0.53 | 0.14 |
| **Sweep rate** | 0.89 | 0.0003 | 0.0001 | 0.06 | 0.007 | 0.02 | 0.08 | 0.04 | 0.02 | 0.8 | 0.005 | 0.67 | 0.46 | 0.3 | 0.07 | 0.28 | 0.03 | 0.46 | 0.7 | 0.22 |
| **P F end** | 0.59 | 0.005 | 0.38 | 0.06 | 0.01 | 0.09 | 0.07 | 0.12 | 0.07 | 0.67 | 0.13 | 0.84 | 0.0006 | 0.016 | 0.0007 | 0.56 | 0.003 | 0.87 | 0.85 | 0.11 |
| **P F max** | 0.75 | 0.68 | 0.0008 | 0.0078 | 0.002 | 0.92 | 0.67 | 0.04 | 0.33 | 0.45 | 0.15 | 0.25 | 0.002 | 0.1 | 0.85 | 0.44 | 0.007 | 0.73 | 0.55 | 0.83 |
| **P F mean** | 0.65 | 0.42 | 0.0006 | 0.008 | 0.41 | 0.66 | 0.56 | 0.005 | 0.89 | 0.2 | 0.52 | 0.19 | 0.11 | 0.49 | 0.13 | 0.62 | 0.89 | 0.54 | 0.6 | 0.54 |
| **BW5** | $<1x10^{-4}$ | 0.79 | 0.66 | 0.01 | 0.24 | 0.46 | 0.44 | 0.43 | 0.21 | 0.34 | 0.88 | 0.01 | 0.72 | 0.22 | 0.44 | 0.54 | 0.77 | 0.2 | 0.2 | 0.51 |
| **BW10** | 0.008 | 0.19 | 0.51 | 0.0006 | 0.0005 | 0.34 | 0.4 | 0.75 | 0.005 | 0.2 | 0.08 | 0.02 | 0.95 | 0.89 | 0.001 | 0.7 | 0.92 | 0.63 | 0.01 | 0.77 |

Supple Table 3 P-values from the statistics comparing calls emitted in the pendulum across different days. Between days, bats vary different call parameters. Each trial of one bat was recorded at a different day. For **F8, F11, F12, M9, M10, M11,** and **M12**, p-values result either from an one-way ANOVA (if data follow a Gaussian distribution based on D´Agostino-Pearson omnibus normality test) or from a Kruskal-Wallis test (if data do not follow a Gaussian distribution). For **F9** and **F10**, p-values result either from an unpaired t-test (if data follow a Gaussian distribution based on D´Agostino-Pearson omnibus normality test) or from a Mann-Whitney test (if data do not follow a Gaussian distribution).

| **Animal ID** | **F1** | **F2** | **F3** | **F5** | **F7** | **M1** | **M3** | **M4** | **M6** | **M7** |
|---|---|---|---|---|---|---|---|---|---|---|
| **N calls/ trial** | 14,13 | 20,21,16 | 15,10,16 | 16,13,16 | 19,20,26 | 12,10,21 | 21,18 | 11,14,11 | 17,18,16 | 13,20 |
| **Interval** | 0.54 | 0.17 | 0.004 | 0.07 | 0.0003 | $<1x10^{-4}$ | 0.73 | 0.65 | 0.09 | 0.25 |
| **Duration** | 0.36 | $<1x10^{-4}$ | $<1x10^{-4}$ | 0.24 | $<1x10^{-4}$ | $<1x10^{-4}$ | $<1x10^{-4}$ | 0.002 | $<1x10^{-4}$ | $<1x10^{-4}$ |
| **Intensity** | 0.002 | $<1x10^{-4}$ | 0.27 | 0.0003 | 0.004 | $<1x10^{-4}$ | 0.11 | $<1x10^{-4}$ | 0.39 | 0.17 |
| **P F start** | 0.002 | 0.78 | 0.005 | 0.0001 | 0.46 | $<1x10^{-4}$ | 0.26 | 0.0005 | $<1x10^{-4}$ | $<1x10^{-4}$ |
| **Sweep rate** | 0.016 | 0.49 | 0.2 | 0.0002 | 0.35 | 0.68 | 0.11 | 0.002 | 0.044 | 0.0001 |
| **P F end** | 0.009 | 0.15 | 0.26 | $<1x10^{-4}$ | 0.46 | $<1x10^{-4}$ | 0.96 | 0.19 | $<1x10^{-4}$ | 0.11 |
| **P F max** | 0.0001 | 0.86 | 0.002 | 0.0004 | 0.66 | 0.0003 | 0.002 | 0.0004 | 0.001 | $<1x10^{-4}$ |
| **P F mean** | 0.0005 | 0.56 | 0.0005 | $<1x10^{-4}$ | 0.46 | 0.0001 | 0.002 | 0.0003 | $<1x10^{-4}$ | 0.0003 |
| **BW5** | 0.032 | 0.0004 | 0.0061 | 0.0004 | $<1x10^{-4}$ | 0.61 | 0.39 | $<1x10^{-4}$ | 0.37 | 0.58 |
| **BW10** | 0.004 | $<1x10^{-4}$ | 0.0053 | 0.25 | $<1x10^{-4}$ | 0.22 | 0.004 | 0.006 | 0.41 | 0.24 |

Supple Table 4 P-values from the statistics comparing changes of call parameters across trials. Red cells represent p-values < 0.05. Blue cells represent p-values > 0.05

| Animal ID (calls/trial) | Trial (Day) | Intensity | Duration | Interval | Sweep rate | P F start | P F end | P F max | P F mean | BW5 | BW10 |
|---|---|---|---|---|---|---|---|---|---|---|---|
| F8 (25, 16, 14, 12, 16, 16, 13, 11) | 1 (1st) | 0.047 | $<1x10^{-4}$ | 0.008 | 0.048 | 0.1 | 0.0014 | 0.7 | 0,144 | 0.002 | 0.0008 |
| | 2 (1st) | 0.73 | 0.51 | 0.07 | >0.99 | 0.77 | >0.99 | 0.02 | 0.08 | 0.13 | 0.52 |
| | 3 (1st) | 0.001 | 0.99 | 0.005 | >0.99 | 0.93 | 0.81 | >0.99 | >0.99 | 0.002 | $<1x10^{-4}$ |
| | 4 (2nd) | 0.13 | >0.99 | 0.8 | 0.034 | >0.99 | 0.036 | 0.95 | >0.99 | >0.99 | 0.63 |
| | 5 (2nd) | 0.72 | >0.99 | 0.33 | >0.99 | 0.37 | >0.99 | >0.99 | >0.99 | >0.99 | 0.73 |
| | 6 (2nd) | 0.23 | >0.99 | 0.036 | 0.19 | >0.99 | 0.12 | >0.99 | >0.99 | >0.99 | 0.33 |
| F9 (28, 15, 10, 15, 17, 23, 18, 34, 16, 16) | 1 (1st) | 0.36 | >0.99 | >0.99 | >0.99 | 0.011 | >0.99 | >0.99 | 0.11 | 0.47 | 0.47 |
| | 2 (1st) | 0.03 | 0.4 | 0.54 | 0.03 | 0.002 | >0.99 | 0.006 | 0.12 | 0.63 | 0.08 |
| | 3 (1st) | >0.99 | 0.58 | >0.99 | 0.22 | 0.029 | 0.46 | 0.003 | 0.02 | >0.99 | >0.99 |
| | 4 (1st) | >0.99 | 0.001 | >0.99 | 0.11 | 0.0076 | >0.99 | $<1x10^{-4}$ | 0.0001 | 0.02 | 0.02 |
| | 5 (1st) | >0.99 | 0.03 | >0.99 | 0.18 | 0.0027 | 0.18 | 0.045 | 0.021 | >0.99 | >0.99 |
| | 6 (2nd) | 0.005 | >0.99 | >0.99 | 0.001 | >0.99 | 0.001 | 0.74 | >0.99 | >0.99 | 0.18 |
| | 7 (2nd) | 0.54 | >0.99 | 0.05 | 0.57 | >0.99 | 0.256 | 0.39 | >0.99 | 0.1 | 0.21 |
| | 8 (2nd) | 0.037 | >0.99 | 0.05 | 0.03 | >0.99 | 0.004 | 0.79 | 0.8 | 0.62 | 0.96 |
| F10 (22, 13, 10, 22, 16, 15, 16) | 1 (1st) | 0.11 | 0.12 | 0.17 | 0.004 | >0.99 | 0.3 | 0.2 | 0.89 | >0.99 | >0.99 |
| | 2 (1st) | 0.08 | 0.01 | 0.38 | 0.009 | >0.99 | >0.99 | 0.002 | 0.17 | >0.99 | >0.99 |
| | 3 (1st) | >0.99 | >0.99 | 0.87 | 0.03 | 0.07 | 0.53 | 0.1 | >0.99 | 0.3 | 0.22 |
| | 4 (1st) | 0.36 | >0.99 | 0.1 | 0.03 | 0.47 | 0.25 | 0.04 | 0.18 | >0.99 | 0.64 |
| | 5 (1st) | >0.99 | >0.99 | 0.67 | 0.22 | >0.99 | 0.05 | >0.99 | 0.71 | >0.99 | >0.99 |
| | 6 (1st) | >0.99 | >0.99 | 0.01 | >0.99 | >0.99 | >0.99 | >0.99 | >0.99 | >0.99 | >0.99 |
| F11 (13, 23, 7, 20, 8, 9, 16, 13, 11, 14, 15, 10, 9, 11) | 1 (1st) | 0.05 | 0.86 | $<1x10^{-4}$ | >0.99 | 0.07 | 0.63 | 0.02 | 0.32 | >0.99 | >0.99 |
| | 2 (1st) | >0.99 | 0.05 | >0.99 | >0.99 | >0.99 | >0.99 | >0.99 | >0.99 | >0.99 | >0.99 |
| | 3 (1st) | 0.65 | 0.56 | 0.04 | >0.99 | 0.61 | 0.04 | 0.006 | 0.006 | 0.73 | 0.71 |
| | 4 (1st) | >0.99 | >0.99 | 0.04 | >0.99 | >0.99 | 0.84 | 0.01 | 0.05 | >0.99 | >0.99 |
| | 5 (2nd) | 0.99 | 0.98 | 0.02 | 0.5 | >0.99 | >0.99 | >0.99 | >0.99 | 0.03 | >0.99 |
| | 6 (2nd) | 0.98 | 0.99 | 0.91 | >0.99 | >0.99 | >0.99 | 0.78 | >0.99 | >0.99 | >0.99 |
| | 7 (2nd) | 0.93 | 0.99 | 0.1 | >0.99 | >0.99 | >0.99 | >0.99 | >0.99 | >0.99 | >0.99 |
| | 8 (2nd) | 0.97 | 0.99 | >0.99 | >0.99 | >0.99 | >0.99 | >0.99 | >0.99 | >0.99 | 0.95 |
| | 9 (2nd) | 0.2 | 0.99 | 0.37 | >0.99 | 0.88 | 0.9 | >0.99 | >0.99 | 0.47 | >0.99 |
| | 10 (2nd) | 0.005 | 0.99 | >0.99 | >0.99 | >0.99 | >0.99 | >0.99 | >0.99 | >0.99 | >0.99 |
| | 11 (2nd) | 0.96 | 0.94 | 0.007 | >0.99 | >0.99 | >0.99 | >0.99 | >0.99 | >0.99 | >0.99 |
| | 12 (2nd) | 0.32 | 0.99 | >0.99 | >0.99 | >0.99 | >0.99 | >0.99 | >0.99 | >0.99 | >0.99 |
| F12 (18, 14, 11, 11, 11, 24, 17, 16) | 1 (1st) | >0.99 | 0.05 | 0.23 | 0.42 | 0.42 | >0.99 | >0.99 | >0.99 | 0.23 | >0.99 |
| | 2 (1st) | >0.99 | 0.55 | 0.001 | 0.04 | 0.07 | >0.99 | 0.17 | 0.49 | 0.38 | 0.04 |
| | 3 (1st) | >0.99 | >0.99 | 0.5 | 0.84 | 0.73 | 0.14- | >0.99 | >0.99 | >0.99 | 0.43 |
| | 4 (2nd) | 0.87 | 0.12 | $<1x10^{-4}$ | $<1x10^{-4}$ | 0.002 | 0.02 | >0.99 | >0.99 | 0.002 | 0.007 |
| | 5 (2nd) | 0.9 | 0.62 | 0.02 | 0.01 | 0.1 | 0.006 | >0.99 | 0.38 | 0.02 | 0.01 |
| | 6 (2nd) | 0.88 | >0.99 | 0.02 | 0.01 | 0.2 | 0.02 | 0.99 | >0.99 | 0.03 | 0.007 |
| M9 (23, 24, 15, 7, 9, 24, 23, 9, 7) | 1 (1st) | 0.008 | 0.005 | >0.99 | 0.04 | $<1x10^{-4}$ | 0.87 | >0.99 | >0.99 | >0.99 | >0.99 |
| | 2 (1st) | >0.99 | 0.07 | 0.03 | >0.99 | 0.01 | 0.92 | 0.47 | >0.99 | 0.88 | 0.41 |
| | 3 (1st) | $<1x10^{-4}$ | 0.46 | >0.99 | >0.99 | 0.07 | 0.25 | 0.07 | 0.07 | 0.0009 | >0.99 |
| | 4 (1st) | >0.99 | >0.99 | 0.94 | 0.006 | $<1x10^{-4}$ | 0.92 | >0.99 | >0.99 | >0.99 | 0.77 |
| | 5 (2nd) | >0.99 | >0.99 | 0.003 | 0.02 | 0.0012 | >0.99 | 0.25 | >0.99 | 0.03 | 0.002 |
| | 6 (2nd) | >0.99 | >0.99 | 0.95 | 0.92 | 0.36 | 0.88 | >0.99 | 0.68 | 0.03 | 0.01 |
| | 7 (2nd) | >0.99 | 0.69 | >0.99 | 0.68 | 0.55 | >0.99 | 0.06 | 0.14 | 0.05 | 0.14 |
| M10 (17, 7, 14, 18, 12, 12, 7, 7) | 1 (1st) | 0.6 | 0.92 | 0.09 | 0.17 | 0.14 | 0.05 | 0.0057 | 0.03 | >0.99 | >0.99 |
| | 2 (1st) | 0.24 | 0.66 | 0.4 | 0.76 | 0.001 | 0.5 | $<1x10^{-4}$ | 0.18 | 0.05 | 0.75 |
| | 3 (2nd) | >0.99 | >0.99 | >0.99 | >0.99 | 0.93 | 0.03 | >0.99 | >0.99 | 0.2 | 0.85 |
| | 4 (2nd) | 0.0001 | 0.0003 | 0.02 | >0.99 | >0.99 | 0.48 | >0.99 | >0.99 | >0.99 | >0.99 |
| | 5 (2nd) | 0.59 | 0.03 | 0.001 | >0.99 | >0.99 | 0.23 | >0.99 | >0.99 | >0.99 | >0.99 |
| | 6 (2nd) | 0.96 | 0.001 | 0.03 | 0.003 | 0.0045 | $<1x10^{-4}$ | >0.99 | >0.99 | 0.6 | 0.73 |
| M11 (22, 8, 12, 11, 10) | 1 (1st) | 0.003 | >0.99 | 0.15 | 0.58 | >0.99 | 0.0008 | 0.09 | 0.003 | >0.99 | 0.9 |
| | 2 (1st) | 0.61 | 0.87 | >0.99 | 0.29 | 0.32 | 0.03 | >0.99 | >0.99 | >0.99 | 0.17 |
| | 3 (1st) | 0.09 | >0.99 | >0.99 | >0.99 | >0.99 | >0.99 | 0.57 | 0.01 | >0.99 | >0.99 |
| | 4 (1st) | 0.0006 | 0.05 | >0.99 | >0.99 | >0.99 | >0.99 | 0.35 | >0.99 | 0.2 | 0.12 |
| | 1 (1st) | 0.86 | 0.88 | 0.22 | 0.26 | 0.94 | 0.049 | 0.12 | >0.99 | >0.99 | >0.99 |
| | 2 (1st) | 0.8 | 0.84 | 0.001 | 0.06 | 0.22 | 0.71 | 0.44 | >0.99 | 0.005 | 0.025 |

| | | | | | | | | | | | |
|---|---|---|---|---|---|---|---|---|---|---|---|
| M12 (20, 10, 7, 11, 13, 11) | 3 (2nd) | 0.28 | 0.27 | >0.99 | 0.07 | 0.96 | 0.14 | 0.67 | 0.97 | >0.99 | 0.97 |
| | 4 (2nd) | 0.67 | 0.61 | >0.99 | 0.009 | 0.16 | 0.95 | 0.23 | 0.94 | >0.99 | 0.97 |
| M13 (15, 29, 11, 24, 20, 20, 14, 16, 12, 15 | 1 (1st) | 0.001 | 0.87 | <1x10$^{-4}$ | 0.0002 | 0.0003 | 0.84 | >0.99 | >0.99 | >0.99 | >0.99 |
| | 2 (1st) | 0.31 | 0.34 | 0.71 | 0.12 | 0.43 | >0.99 | 0.7 | >0.99 | >0.99 | >0.99 |
| | 3 (1st) | >0.99 | 0.1 | 0.05 | 0.0008 | <1x10$^{-4}$ | >0.99 | >0.99 | 0.96 | >0.99 | >0.99 |
| | 4 (2nd) | >0.99 | >0.99 | 0.55 | 0.002 | 0.0002 | 0.85 | 0.33 | 0.14 | 0.006 | >0.99 |
| | 5 (2nd) | >0.99 | >0.99 | 0.05 | 0.004 | 0.0015 | 0.8 | 0.19 | 0.07 | 0.08 | >0.99 |
| | 6 (2nd) | >0.99 | >0.99 | >0.99 | 0.35 | 0.12 | >0.99 | 0.2 | 0.17 | 0.06 | 0.01 |
| | 7 (2nd) | 0.08 | <1x10$^{-4}$ | 0.02 | >0.99 | >0.99 | 0.0001 | 0.07 | 0.08 | >0.99 | 0.09 |
| | 8 (2nd) | >0.99 | >0.99 | >0.99 | 0.03 | 0.11 | 0.01 | 0.63 | 0.1 | 0.04 | 0.21 |
